# STAR Suite: Transcriptomics processing in a single binary through AI-assisted development

**DOI:** 10.64898/2026.03.09.710580

**Authors:** Ling-Hong Hung, Dylan Baker, Bill Flynn, Danwei Huangfu, Renhe Luo, Paul Robson, Ting Zhou, Ka Yee Yeung

## Abstract

**Background:** STAR is the aligner underlying most transcriptomics processing, and is used inside Cell Ranger. However, the surrounding steps—adapter trimming, sorting, feature-barcode assignment, probe handling, SLAM-seq analysis, quantification, and quality control—run as separate tools scripted around it. Cell Ranger hides this complexity but is proprietary, restricted to 10x products, and barred by its license from redistribution, so it cannot serve as a shared processing layer. While STARsolo supports scRNA-seq, it lacks the feature-barcode detection needed for Perturb-seq, where short tags such as CRISPR guides or lineage barcodes are read alongside the transcriptome. No open-source tool processes Perturb-seq feature barcodes at production scale, and the only open-source processor for 10x Flex—a probe-based assay for fixed and FFPE tissue that multiplexes several samples in one lane—is a recent standalone k-mer tool.

**Results:** STAR Suite compiles these steps into the STAR codebase to create a drop-in executable for bulk RNA-seq, scRNA-seq, Perturb-seq, 10x Flex, and SLAM-seq. It adds no dependencies beyond OpenSSL’s libcrypto, and legacy behavior is preserved. On consortium and public benchmarks it is up to 41-fold faster than Cell Ranger 9.0.1 for Flex (20- to 41-fold from binary CBQ input, 17- to 23-fold from FASTQ) and 3.9- to 6.2-fold faster for scRNA-seq and Perturb-seq, timed on one modest server (24 cores, 128 GiB, local SSD) except for the largest dataset, which ran on a cloud instance limited to the same size. The concordance to Cell Ranger is high: gene-level Spearman and Pearson 0.98–1.0, cell-level Pearson on per-cell total counts 0.9999–1.0, feature-UMI Pearson 0.999–1.000, Jaccard agreement of the called-cell sets 0.97–0.995, and 99–100% CRISPR-call agreement. It also needs far less disk: its peak use on the largest Flex dataset is 16.9 GiB, against 1.8 TiB for Cell Ranger. These gains arise from running the steps in one process rather than as separate programs exchanging files, and from four new computational techniques: fast-Hamming, a vectorized exact Hamming-distance search, used for Flex probe matching and optionally for feature barcodes; a disambiguated hash cascade that assigns Flex probes without alignment; a permit-based thread scheduler that interleaves alignment and feature assignment; and variance-based auto-trimming for SLAM-seq conversion calling, which finds the reliable region of each read from how variable the conversion rate is along it. The four modules add 183,606 lines of C/C++ to STAR’s 28,228. STAR Suite is the NIH MorPhiC consortium’s production processor. The same binary is run by people, by AI agents and by cluster schedulers, and every production run is recorded with its commands, checksums and outputs. It has been built and maintained over eight months and 25 releases under a human-directed, AI-implemented workflow, with its full design and benchmark record public.

**Conclusions:** STAR Suite provides the first production-ready open-source implementation of Perturb-seq feature-barcode processing, and an open end-to-end 10x Flex pipeline matched to Cell Ranger, delivered as a drop-in replacement for the STAR aligner. The design also extends: downstream analysis now done in Python or R packages can be built into the same binary, and further modalities added alongside, as a companion preprint does for ATAC-seq. Source code, workflow recipes, and per-run provenance are released under the MIT license.

## 1 Background

Processing the raw sequencing data from today’s transcriptomic assays—bulk RNA-seq, single-cell RNA-seq (scRNA-seq)^1^, Perturb-seq^2,3^, 10x Genomics Flex (fixed RNA profiling) ^4^, and metabolic labeling methods such as SLAM-seq ^5,6^—means running a multi-stage pipeline: separate tools for demultiplexing, barcode parsing, sequence alignment, deduplication, and feature counting, each with its own inputs, outputs, and dependencies. The analysis that follows the counts—clustering, differential testing, model fitting—is a further layer, usually of Python and R packages. At the core of most of these pipelines lies the open-source Spliced Transcripts Alignment to a Reference (STAR) aligner^7^, which STARsolo ^8^ extends to single-cell gene counting and which users now commonly run under a community-tuned parameter set. Everything around the aligner—adapter trimming, sample-aware execution, feature-barcode assignment, probe handling, SLAM-seq analysis, sorting, quantification, and quality control (QC) reporting—is still a separate tool that users script around STAR.

A researcher processing this data has two practical options. The first is to chain specialized open-source tools through a workflow language, the standard route to scalable, reproducible processing. Nextflow ^9^, with its nf-core ^10^ community pipelines, comes closest to general adoption among the workflow languages (WDL ^11,12^ and CWL ^13,14^ are the others in common use), and graphical builders such as our own Biodepot-workflow-builder (Bwb) remove most of the hand-written scripting ^15–18^. These systems are reliable, but the pipeline underneath is still a set of separate programs: every stage writes and re-reads intermediate files, which on a large single-cell run can mean terabytes of scratch and can decide where the run is possible at all. Each program carries its own library stack, and keeping the whole installable and reproducible means containers that must themselves be maintained. That remains too complex for the many researchers who do not write and maintain pipelines themselves.

The second option is an integrated platform. Cell Ranger, the de facto standard for 10x single-cell data, folds the same steps into one simple command through an internal orchestration language (Martian), so the bench user never handles the orchestration. However, Cell Ranger is not only proprietary; its end-user license also prohibits redistribution and sublicensing, disallows modification or derivative works, and restricts use to data generated with 10x products. Those terms are manageable for a laboratory analyzing its own 10x data; they rule Cell Ranger out as a shared, redistributable processing layer.

This work was built for the NIH Molecular Phenotypes of Null Alleles in Cells (MorPhiC) consortium ^19^, which systematically knocks out every human protein-coding gene and profiles the resulting molecular phenotypes. Its requirements sharpen the problem. The consortium’s processed data is a public resource: every run must be reproducible and auditable, the methods transparent enough for the community to vet, and the procedures easily applied to a user’s own data so that comparisons with MorPhiC results are possible. That calls for one open processor, run the same way for all of the consortium’s assays—bulk RNA-seq, scRNA-seq, Perturb-seq, multiplexed Flex, and SLAM-seq. Moreover, to meet the FAIR principles ^20^—findable, accessible, interoperable and reusable—now standard for shared data, every released result should be traceable to the process that produced it: the inputs, outputs, commands, scripts and executables, with their versions and dependencies. Finally, the consortium’s data coordination center now uses AI agents to write and run much of its production processing, sharpening the need for procedures that can be reproducibly composed by agents and verified by humans.

Neither alternative meets these requirements: Cell Ranger’s terms rule it out, and orchestrating separate tools makes auditing and reproducible agentic use difficult, quite apart from the time, memory and disk that each run demands. In addition, there were gaps in the tool coverage. No open-source tool could process feature barcodes at production scale, which left Cell Ranger as the only realistic option. For Flex, the one open-source processor is a recent standalone k-mer tool, scalable but calling cells on each tag separately rather than on the sample, which produces significant differences in cell calling relative to Cell Ranger on samples with multiple tags (Results). The obvious remedy, creating the missing tools and integrating the surrounding steps into STAR itself, had not been taken because it is far harder than scripting around the aligner. Even adding the simple ability to run several samples through STAR in one invocation, rather than reloading the genome index for each, means threading new control flow through tens of thousands of lines of carefully optimized C/C++ without changing anything else. For a small academic group that has been infeasible, which is why even 10x Genomics wrapped an unmodified STAR in its own pipelines rather than modify the aligner. Two things have changed. The surrounding methods have matured— alignment, cell calling, adapter trimming, sorting and quantification are now standard enough that compiling them into one binary loses little flexibility—and human-directed AI software engineering has made modification at this depth practical. With an architectural plan, gold-standard tests taken from production data, and AI agents writing the implementation, a single developer-scientist can make focused changes to the STAR code and robustly update and maintain the new codebase.

### 1.1 Our contributions

We built STAR Suite, a drop-in, open-source replacement for the STAR executable that processes bulk RNA-seq, scRNA-seq, Perturb-seq, 10x Flex, and SLAM-seq from FASTQ—or, experimentally, from the binary CBQ format of BINSEQ^21^—to standard count outputs (Fig. 1). It adds no dependencies beyond OpenSSL’s libcrypto and retains every function of the original aligner, so anyone already running STAR can adopt it without changing how they work. For a researcher, each assay is a single command—no workflow language, no container, no wrapper scripts, and nothing new to install. The new functions are enabled by flags, and the executable composes with upstream and downstream tools through scripts or workflow languages. It brings established methods into the binary: Cutadapt-compatible adapter trimming, memory-bounded BAM sorting, Y-chromosome removal, Variational Bayes (VB) / expectation–maximization (EM) transcript quantification (TranscriptVB), SLAM-seq analysis with per-sample variant masking, and parallel reading of the blocked-gzip FASTQ that sequencers produce. It introduces four new computational techniques: fast-Hamming, a vectorized exact Hamming-distance search that decides Flex probe matches and, where a hash cannot cover the required distance, feature-barcode matches; a disambiguated hash cascade that assigns Flex probes without alignment; a permit-based thread scheduler that interleaves alignment and feature assignment, enabled by the single-process integration; and variance-based auto-trimming for SLAM-seq.

**Figure 1:**
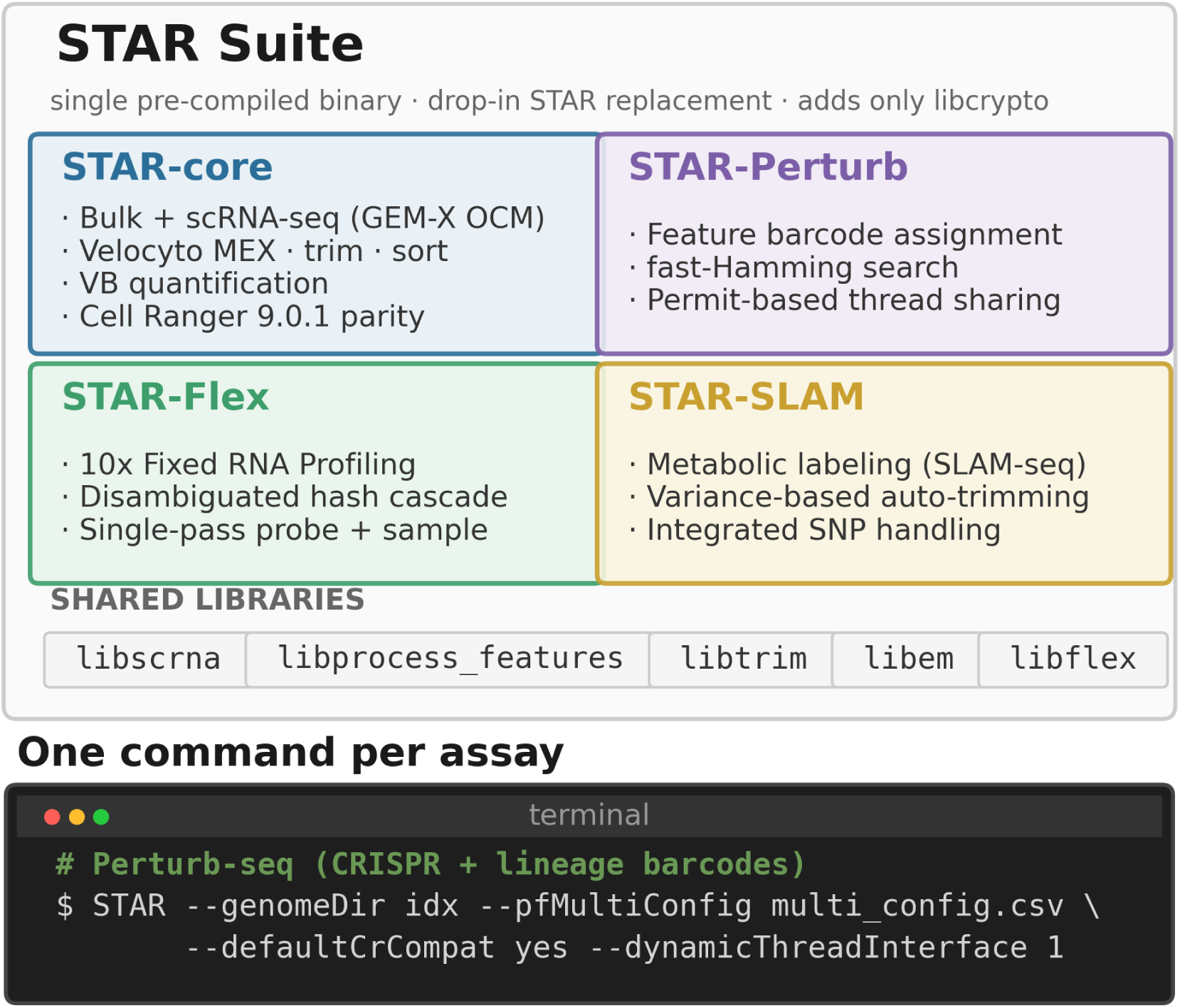
STAR Suite: architecture and single-command interface. Four processing modules—STAR-core (bulk and single-cell RNA-seq), STAR-Perturb (feature-barcode assays), STAR-Flex (probe-based Flex), and STAR-SLAM (metabolic labeling)—share five C/C++ libraries (`libscrna`, `libprocess_features`, `libtrim`, `libem`, `libflex`) inside a single binary. Adapter trimming, transcript quantification, feature-barcode assignment, Flex probe processing, SLAM-seq analysis, BAM sorting, Y-chromosome removal, and QC are integrated natively, and each assay is invoked through the standard STAR executable with a small set of module-specific flags. All legacy STAR functionality is preserved when the new flags are not set.

These modules close the two gaps noted above: `process_features`, used by STAR-Perturb, is the first open-source implementation of feature-barcode processing at production scale, and STAR-Flex reproduces Cell Ranger’s multiplexed Flex output—sample demultiplexing, per-sample cell calling and count matrices—end to end. Both these implementations began as separate executables scripted around the aligner, but the integrated versions proved simpler, faster and closer to Cell Ranger (Results). Because Cell Ranger’s outputs are what the field has built on, close agreement with them is a requirement rather than a courtesy: a user who switches should reach the same biological conclusions. STAR Suite was built with AI assistance—183,606 lines of new C/C++ around STAR’s 28,228—and has been maintained in production for eight months and 25 releases, with every design document and benchmark record kept in the public repository (Results). Because the whole pipeline for an assay is one command, a script template (recipe) and a run record can describe processing completely, which is what the consortium’s traceability requirement needs. The sections that follow benchmark the four modules against Cell Ranger 9.0.1 and the stepwise pipelines they replace, then turn to the development record and to the interface through which people and agents run the suite.

## 2 Results

STAR Suite is organized as four modules within a single binary (Fig. 1). STAR-core provides the shared alignment, counting, and QC routines that underlie all modules, handling both bulk and single-cell RNA-seq. STAR-Perturb, STAR-Flex, and STAR-SLAM each extend this core with assay-specific processing—feature barcode assignment, probe-based quantification, and metabolic-labeling analysis, respectively—activated by a small number of module-specific flags. Related flags are grouped into preset modes (for example, one flag applies the Cell Ranger-compatible defaults of the Perturb-seq path), and the repository provides worked examples for every assay type (see Methods).

We benchmarked the four modules of STAR Suite using data generated by the MorPhiC program and public data (see Methods and Table 1). The consortium data are the production datasets STAR Suite processes: the MSK 30-KO Perturb-seq dataset and the JAX SC2300771 Flex dataset. Because those datasets remain under consortium embargo, every assay also has a public benchmark, so the comparisons reported here can be rerun independently. Each public dataset was chosen to provide a set of representative benchmarks (Table 1): the 10x PBMC 10K set is the standard 3’ gene-expression reference, distributed with Cell Ranger’s own outputs; the A375 set is 10x’s public Perturb-seq dataset, with its guide calls; GSE325982 is a Flex dataset whose submitters deposited their own Cell Ranger 9.0.1 outputs, an independent check on the reference run we made for it; the 320k scFFPE set brings scale (7.3 billion read pairs, 16 sample tags) and the newer GEM-X chip generation; and the GRAND-SLAM fixture is the reference implementation’s own example data.

**Table 1.** Benchmarking datasets. Tags are 10x sample barcodes and lanes are sequencing lanes. Cell counts are the reference tool’s, and for A375 the count is of cells carrying a guide call rather than of called cells. The MSK 30-KO and JAX SC2300771 datasets are MorPhiC consortium data under embargo (Availability of data and materials); the rest are public. GEX, gene expression; PE, paired end.

| Dataset | Assay | Scale | Availability |
| --- | --- | --- | --- |
| Paired-end bulk (PPARG revertant WT replicate 3, KOLF2.2J iPSC) | Bulk RNA-seq | 35.1M paired end (PE) reads | GEO GSE288287; ENA https://www.ebi.ac.uk/ena/browser/view/ERP166966 |
| 10x PBMC 10K (3' v3 chemistry) | scRNA-seq | 638.9M PE reads; 11,806 cells (Cell Ranger) | 10x Genomics dataset [10-k-pbm-cs-from-a-healthy-donor-v-3-chemistry-3-standard-3-0-0](https://www.10xgenomics.com/datasets/10-k-pbm-cs-from-a-healthy-donor-v-3-chemistry-3-standard-3-0-0) |
| A375 CRISPR 5' GEX | Perturb-seq | 11 features; 1,083 cells carry a Cell Ranger guide call | https://www.10xgenomics.com/datasets/1k-CRISPR-5p-gemx |
| MSK 30-KO (ES sample) | Perturb-seq | 33,226 cells; 30 CRISPR guides + 245,979 LARRY barcodes | MorPhiC consortium release (pending) |
| JAX SC2300771 | 10x Flex | 4 tags, 8 lanes; 2.011B PE reads | ENA ERP189111 |
| GSE325982 Flex pool (4 single-tag samples) | 10x Flex | 4 tags, 1 lane; 1.121B PE reads | GEO GSM9618375; ENA SRX32655594 |
| 10x 320k scFFPE (8 human tissues) | 10x Flex (GEM-X, 16-plex) | 16 tags, 4 lanes; 7.3B PE reads | https://www.10xgenomics.com/datasets/320k_scFFPE_16-plex_GEM-X_FLEX |
| GRAND-SLAM 100K human | SLAM-seq | 100K reads | https://github.com/erhard-lab/gedi/wiki/GRAND-SLAM |

We evaluated STAR Suite on two criteria: agreement of the outputs with the reference, and performance—runtime, memory and, for Flex, disk use—at production scale. Tables 2 and 3 give the outcome first, speed and concordance on every benchmark; the subsections that follow describe the shared core and then explain, module by module, where each gain comes from and where each residual difference localizes.

**Table 2.** Wall-clock time and speedup of STAR Suite 1.9.5 against the reference pipeline on each benchmark. The reference is Cell Ranger 9.0.1—no BAM output, secondary analysis disabled, timed to completion—except for bulk RNA-seq, whose reference is an external stepwise pipeline (Trim Galore + STAR + Salmon in alignment mode with automatic library-type detection; the with-Y-removal reference adds an external `awk`/`samtools`/`gzip` Y-removal step). SLAM-seq has no row: its reference pipeline does different work (Results). The MSK 30-KO row processes gene expression, CRISPR guides and the LARRY lineage library in one STAR Suite pass against the sum of the two Cell Ranger runs that are needed to obtain the same results. All rows except the 320k dataset were measured on one machine (Intel i9-13900KF, 24 cores, 32 threads) with every file on local NVMe, for STAR Suite and the reference alike; the 320k rows were measured for both tools on an AWS instance of the same size with local NVMe (Methods). CBQ rows read binary input converted once from the FASTQ (conversion excluded); every other row reads the original FASTQ. Of the rows that read FASTQ, the PBMC, A375 and original-BGZF Flex rows are read by the in-process parallel BGZF reader; the MSK files are ordinary gzip and were read through STAR’s internal gzip route, and the bulk runs use STAR’s external `zcat` route, as their production commands do. For the modules that align, the route makes little difference to the time (Results). Exact seconds are in Supplementary Table S5; the spinning-disk comparison is in Supplementary Table S9.

| Assay | Benchmark (input) | STAR Suite | Reference | Speedup |
| --- | --- | --- | --- | --- |
| Bulk RNA-seq | PPARG 35.1M PE, no Y-removal | 8 min 6 s | 12 min 49 s | 1.6× |
| Bulk RNA-seq | PPARG 35.1M PE, with Y-removal | 9 min 35 s | 40 min 47 s | 4.3× |
| scRNA-seq | 10x PBMC 10K, 3’ gene expression (639M pairs) | 16 min 18 s | 63 min 0 s | 3.9× |
| Perturb-seq | A375, gene expression + CRISPR guides | 2 min 26 s | 11 min 25 s | 4.7× |
| Perturb-seq | MSK 30-KO ES, gene expression + guides + LARRY, one pass | 27 min 6 s | 167 min 2 s | 6.2× |
| 10x Flex | JAX SC2300771 (2.01B pairs), original BGZF FASTQ | 3 min 20 s | 57 min 6 s | 17.2× |
| 10x Flex | JAX SC2300771, CBQ | 2 min 50 s | 57 min 6 s | 20.2× |
| 10x Flex | GSE325982 (1.12B pairs), plain gzip FASTQ via rapidgzip | 4 min 17 s | 74 min 6 s | 17.3× |
| 10x Flex | GSE325982, CBQ | 1 min 48 s | 74 min 6 s | 41.0× |
| 10x Flex | 10x 320k scFFPE (7.30B pairs), original BGZF FASTQ | 17 min 36 s | 6 h 37 min 38 s | 22.6× |
| 10x Flex | 10x 320k scFFPE, CBQ | 10 min 45 s | 6 h 37 min 38 s | 37.0× |

**Table 3.** Concordance between STAR Suite 1.9.5 and the reference tool on the benchmarks in Table 1. Barcode Jaccard is the fraction of all called cells that both tools called. Gene Spearman and gene Pearson are on raw per-gene totals over the cells both tools called, over every gene in both annotations. Cell Pearson is on each shared cell’s total UMI count, across cells; mean per-cell Pearson correlates each shared cell’s log-transformed counts across genes with at least 20 counts in both outputs and detected in at least 1% of shared cells, averaged over cells (Methods). The multiplexed Flex rows pool all samples, with a cell identified by its barcode and sample tag; per-sample values are in Supplementary Table S3. Feature-UMI Pearson is on per-guide UMI sums over shared cells, and CRISPR calls is the fraction of shared cells whose called guide set is identical. NTR, new-to-total ratio. Residual-discrepancy sources are localized in Table 5; further detail is in Supplementary Tables S1–S4.

| Assay | Benchmark (reference) | Concordance |
| --- | --- | --- |
| Bulk RNA-seq | PPARG (Salmon) | gene Spearman 0.9992, Pearson 1.0000; transcript read-count Spearman 0.985, Pearson 0.99998 |
| scRNA-seq | 10x PBMC 10K (Cell Ranger 9.0.1) | barcode Jaccard 0.995; cell Pearson 0.99999; mean per-cell Pearson 0.992; gene Spearman 0.990, Pearson 0.9998 |
| Perturb-seq | A375 (Cell Ranger 9.0.1) | CRISPR calls 100% (1,076 shared cells); feature-UMI Pearson 0.99999; barcode Jaccard 0.992; cell Pearson 0.99995; mean per-cell Pearson 0.992; gene Spearman 0.988, Pearson 0.980 |
| Perturb-seq | MSK 30-KO ES (Cell Ranger 9.0.1) | CRISPR calls 99.0%; feature-UMI Pearson 0.9994; barcode Jaccard 0.992; cell Pearson 0.99998; mean per-cell Pearson 0.991; gene Spearman 0.993, Pearson 0.9994 |
| 10x Flex | JAX SC2300771 (Cell Ranger 9.0.1) | barcode Jaccard 0.988; cell Pearson 0.99999; mean per-cell Pearson 0.9986; gene Spearman 0.99997, Pearson 0.9999997 |
| 10x Flex | GSE325982 (Cell Ranger 9.0.1) | barcode Jaccard 0.994; cell Pearson 0.99997; mean per-cell Pearson 0.9952; gene Spearman 0.99996, Pearson 0.9999993 |
| 10x Flex | 10x 320k scFFPE (Cell Ranger 9.0.1) | barcode Jaccard 0.974 (333,411 STAR cells against 325,410); cell Pearson 0.99998; mean per-cell Pearson 0.9990; gene Spearman 0.9999997, Pearson 0.9999997 |
| SLAM-seq | GRAND-SLAM 100K (GRAND-SLAM) | NTR Pearson 0.9989 / 0.9961 / 0.9944 at $\geq 20$ / 50 / 100 reads |

For scRNA-seq, Perturb-seq, and Flex, STAR Suite produces Cell Ranger-compatible Market Exchange (MEX) matrices and assay-specific auxiliary files (CRISPR call tables, sample-tag assignments); for bulk RNA-seq and SLAM-seq, it produces the standard count and new-to-total RNA ratio (NTR) outputs used by downstream R (Seurat^22^, Bioconductor/DESeq2^23,24^) and Python (Scanpy/AnnData ^25,26^) analysis ecosystems. STAR Suite performs this end-to-end processing within a single process, instead of the orchestrated, multi-stage pipeline Cell Ranger runs. This is what makes the permit-based thread scheduling described below possible and lets Flex probe assignment, sample-tag detection and cell calling run in a single pass rather than the separate passes a standalone tool requires. For bulk RNA-seq, the comparison is instead with a conventional stepwise pipeline, whose adapter trimming, sorting, and transcript quantification STAR Suite folds into the same single invocation.

The concordance measures in Table 3 compare the two tools’ outputs for genes, for cells and for the called-cell set. Gene Spearman and gene Pearson are computed on each gene’s count summed over the cells both tools called: one total per gene for each tool, correlated across genes, which asks whether the tools assign reads to the same genes in the same proportions. Cell Pearson correlates each shared cell’s total unique molecular identifier (UMI) count between the tools, across cells, which asks whether each cell receives the same number of molecules. Mean per-cell Pearson is a different quantity: for each shared cell, the two tools’ counts are correlated across genes, and these correlations are averaged over cells, which asks whether each cell has the same expression profile. Barcode Jaccard is the number of cells both tools call divided by the number either tool calls, so a Jaccard of 0.995 means that 0.5% of the cells called by either tool were called by only one of them.

Datasets that multiplex several samples, each read carrying a sample tag, can be compared in two ways. Pooling all samples and computing each measure once ignores the samples, so the comparison reflects cell calling and counting. Comparing each sample separately and averaging the results also brings in any disagreement in assigning reads to samples, which shifts counts between samples without changing their pooled totals. In both, a cell is identified by its barcode together with its sample tag, because cells from different samples can share a barcode. The values quoted in the abstract and in Table 3 pool the samples and treat them as one; the per-sample values are given with the Flex results and in Supplementary Table S3.

### 2.1 STAR-core: input, sorting, cell calling and threads shared by every module

Several of the changes in STAR Suite sit in the shared core rather than in one assay’s pipeline, and every module that uses them inherits them. Table 4 lists every change, shared and module-specific, with the modules it applies to. We describe the shared ones once here; the assay subsections that follow build on them.

**Table 4.**
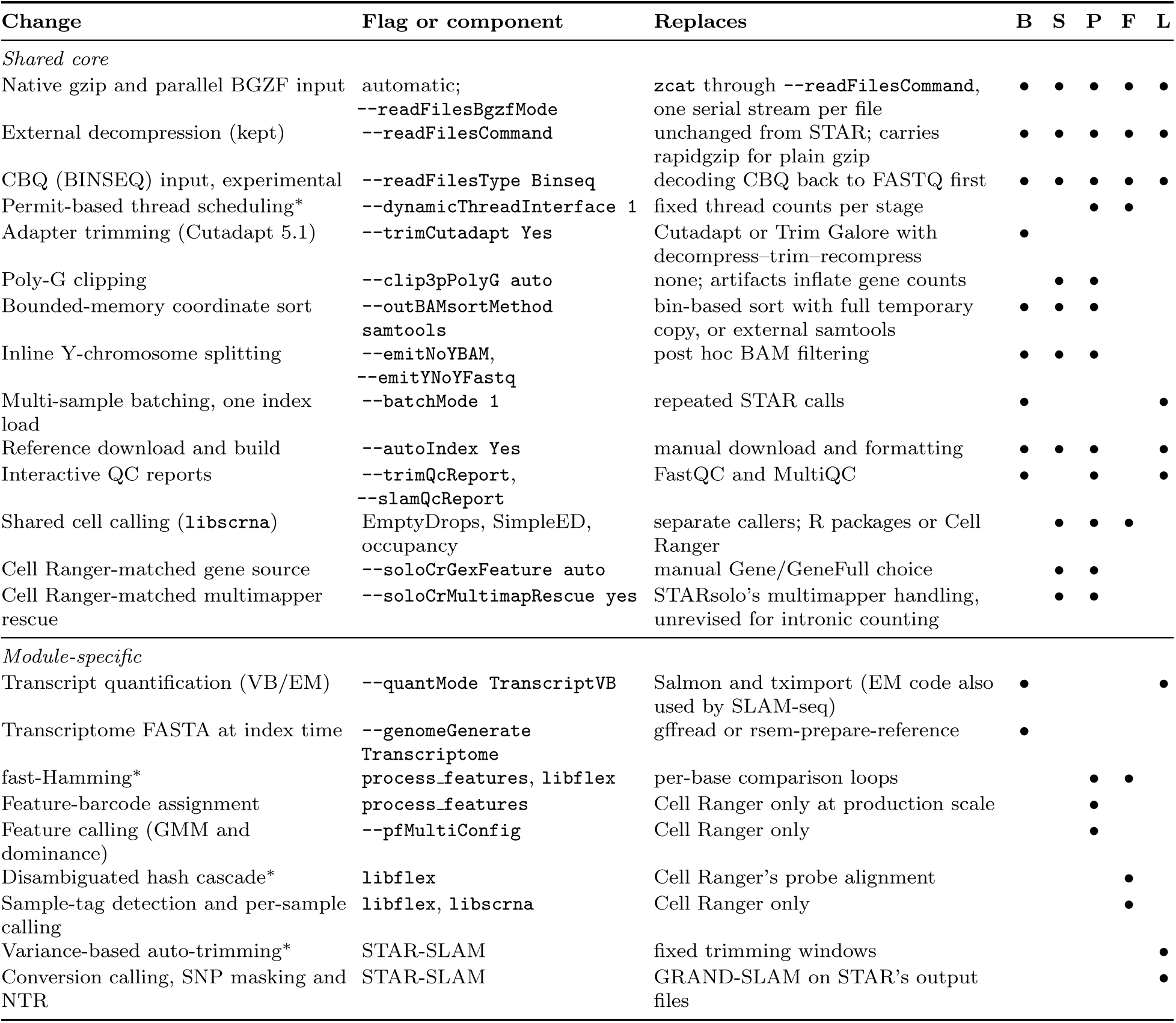
What STAR Suite changes and where each change applies. Bullets mark where a change is used routinely, not every module it is available to: adapter trimming runs in any aligned run but is normally needed only for bulk RNA-seq, and any run that writes coordinate-sorted BAM uses the bounded-memory sorter. B, bulk RNA-seq; S, scRNA-seq; P, Perturb-seq; F, Flex; L, SLAM-seq. Changes marked *^∗^* are new techniques introduced in this work, each described in its Methods subsection. Parallel BGZF input is not used while TranscriptVB learns its model online, and the external decompression route is STAR’s own, kept unchanged.

Upstream STAR reads compressed FASTQ through an external command, typically `zcat`, as one serial stream per file. STAR Suite has an internal gzip reader, used by all modules. It also detects BGZF—the blocked gzip (bgzip) format Illumina sequencers use for FASTQ—and decompresses its blocks in parallel inside the process (Fig. 2; Methods). The external route is retained for compatibility and to allow use of a helper application such as rapidgzip to parallelize decompression of ordinary gzip FASTQ files. For the modules that align, reading is a small part of the run and the gain is correspondingly small. For STAR-Flex, which aligns nothing, the speed of reading the input bounds the whole run (Table 2). The GSE325982 files are distributed by the archive as ordinary gzip; read through that external route with rapidgzip they took 4 min 17 s, against 9 min 7 s through STAR’s internal gzip reader. Besides FASTQ, CBQ binary input is also supported experimentally. A new bounded-memory backend for coordinate sorting, `--outBAMsortMethod samtools`, is modeled on `samtools sort` and replaces STAR’s bin-based sorter. The bin-based sorter writes the whole alignment set to temporary files and needs enough memory for its fullest bin; the new backend holds records in memory up to a set limit and writes sorted runs to disk only beyond it (Methods). It is used by any run that writes coordinate-sorted BAM, and its SAM records are identical to those of samtools sort (Table 5).

**Figure 2:**
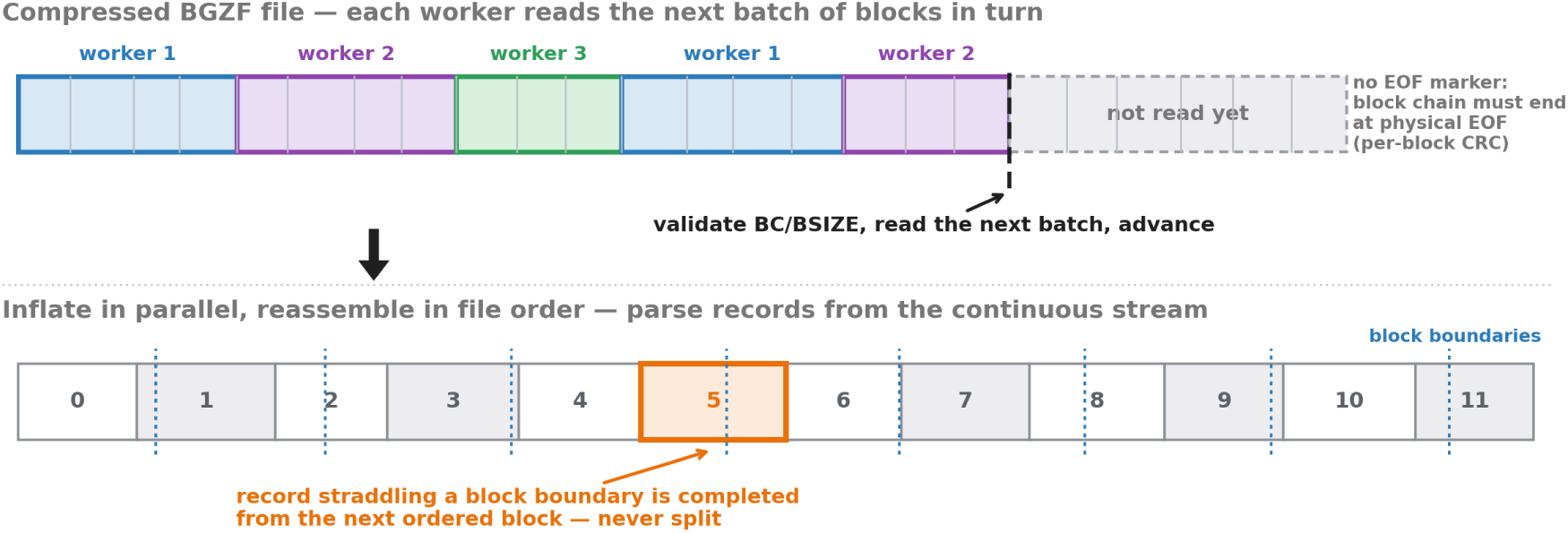
Parallel ingestion of sequencer-original BGZF FASTQ. Top: each inflate worker reads the next run of compressed blocks, bounded in size. The end of the file and every block’s CRC is checked. Bottom: batches are inflated in parallel, reassembled in file order and parsed as one stream, so a record that straddles a boundary is completed rather than split. Records are handed to the processing threads in batches (Methods).

**Table 5.** Parity targets and residual-discrepancy localization across STAR Suite components. Bracketed numbers index the variance classes defined in Supplementary Table S6. Rows without a bracketed number have no counterpart among those classes. CR, Cell Ranger.

| Component | Reference target | Output evaluated | Residual-discrepancy source | Interpretation for intended use |
| --- | --- | --- | --- | --- |
| Adapter trimming | Cutadapt / Trim Galore | Trimmed FASTQ output | Cutadapt version differences: default v5.1, with `--trimCutadaptCompat` Cutadapt3 for legacy v3.2 parity [10] | Functional byte equivalence against the selected Cutadapt version |
| Coordinate sorting | External **samtools sort** | Coordinate-sorted BAM (SAM-record content) | BGZF block boundaries not deterministic across builds; SAM records identical | Functional byte equivalence at the SAM-record level |
| QC summaries | FastQC | Integrated trim-QC metrics | Report-format and presentation differences (primary-only vs all-records) | Same QC interpretation without an external FastQC and MultiQC dependency |
| Reference construction | Cell Ranger references / probe references | Compatible STAR + Flex references | Index binaries not byte-identical across STAR builds even with identical FASTA and GTF; naming, gene filtering, annotation choices | Documented source-asset reconstruction; index variation is downstream-tolerant |
| TranscriptVB | Salmon alignment-mode + tximport | Gene counts (production); transcript estimates (diagnostic) | Per-read processing order under multithreading [5]; partial sampling state in Salmon dynamic auto-detection [4]; inherent isoform-level ambiguity in the transcript estimates [9] | Gene Spearman 0.999, Pearson 1.000; transcript read-count Spearman 0.985, Pearson 0.99998; single-threaded matched-order reproduces Salmon |
| Cell filtering | EmptyDrops / Cell Ranger-style **libscrna** | Called cells + cell-level expression | Marginal barcode ranks [2]; EmptyDrops Monte Carlo boundaries and RNG state [1] | High-confidence cell population stable; exact marginal calls are not the biological target |
| scRNA-seq processing | Cell Ranger 9.0.1 | GEX matrices + cell calls | Tie resolution in UMI deduplication and multimapper assignment [3]; intronic and lncRNA regions [7]; overlapping and poorly annotated genes [8]; CR7 intronic-counting default [10] | Gene Spearman 0.990, Pearson 0.9998; Jaccard 0.995; cell Pearson 0.99999, mean per-cell Pearson 0.992; the remaining gap to upstream STARsolo is not closable by parameter tuning |
| Feature processing (Perturb) | Cell Ranger feature-barcode outputs | Feature UMI counts + guide/lineage calls | Listed feature matches present in input but absent in CR (clean-room confirmed); connected-component merging vs CR's documented 1-bp UMI correction<sup>38</sup> [12]; EmptyDrops Monte Carlo [1] | 99–100% CRISPR match; UMI Pearson 0.999–1.000; missing CR counts not pursued as design target |
| Flex processing | Cell Ranger Flex 9.0.1 | Sample tags + cells + gene matrix | Naive and offset-tolerant split-probe matching do not reproduce the implementation [11]; strict fixed-offset matching does once tag ties [3], marginal cells [2] and pooled-tag cell calling [2] are matched; unique-single-mismatch tag rule established against BAM output | Jaccard 0.988 (JAX), 0.994 (GSE325982), 0.974 (320k); gene Spearman and Pearson both above 0.9999; cell Pearson above 0.9999, mean per-cell Pearson 0.995–0.999 |
| SLAM-seq NTR | GRAND-SLAM / GEDI compatibility | Gene-level NTR and k/nT | Overlap rules, SNP masks, trim windows; GEDI compatibility choices; digamma-function implementation differences in EM [6] | NTR Pearson 0.994–0.999 across read-count thresholds; compatibility mode isolates reference-specific behavior |

Adapter trimming reimplements Cutadapt within the aligner, so a trimmed run no longer decompresses, trims and recompresses its FASTQ, and its output is byte-identical to that of the selected Cutadapt version. Y-chromosome reads can be split out while bulk and single-cell alignments are written, the step the consortium’s cell line requires (Bulk RNA-seq, below).

Our updated cell-calling library, `libscrna`, is used by scRNA-seq, Perturb-seq and Flex (Methods), so improvements to it reach all three. Two key changes were made in the 1.9 releases: sorting each bootstrap sample’s barcode counts once and sharing the thread budget across samples cut the cell-calling step on one 1.8-billion-read-pair lane of the 320k dataset from 244 to 18 s, and spreading the Monte Carlo draws over eight threads cut the A375 caller from 24.0 to 3.7 s.

The threads of a run are allocated by the permit-based thread scheduler (Methods): alignment, feature assignment and, in STAR-Flex, block decompression and cell calling draw on one thread budget within the single process, so a stage that is waiting does not hold threads another stage could use. This currently applies to STAR-Flex and STAR-Perturb.

### 2.2 Bulk RNA-seq: the base production application

Bulk RNA-seq is the most widely used transcriptomics assay and was the entry point that motivated the entire STAR Suite integration effort. One requirement is particular to the consortium. The KOLF2.2J iPSC line used for most MorPhiC experiments, a derivative of the KOLF2.1J reference line ^27^, is licensed under terms that require Y-chromosome reads to be removed before sequence data is redistributed, so the shared pipeline strips them routinely. In production-scale consortium processing on the previous fragmented toolchain, the pre- and post-alignment steps—adapter trimming, BAM sorting, Y-chromosome removal, transcriptome FASTA generation—routinely took longer than the alignment itself, with a substantially larger intermediate-file disk footprint. STAR-core internalizes these stages into a single STAR invocation: adapter trimming, inline Y-chromosome splitting and bounded-memory sorting as described above, together with three additions this assay needs. Transcriptome BAM generation and Variational Bayes / EM transcript quantification (--quantMode TranscriptVB) replace the external Salmon and tximport step. Native multi-sample execution loads the genome index once per run, rather than relying on repeated STAR calls or on shared-memory caching that may fail in containerized environments. HTML and JSON QC reports remove the external FastQC and MultiQC dependency. Table 4 summarizes the additions and the legacy workarounds each replaces.

On the paired-end PPARG benchmark (35.1M PE reads, KOLF2.2J iPSC; Table 1), the integrated STAR Suite runs 1.6-fold faster than an external Trim Galore + STAR + Salmon pipeline without Y-chromosome removal and 4.3-fold faster with Y-removal (Table 2). In the consortium, most experiments—and all bulk RNA-seq to date—use the KOLF2.2J line, which requires Y-chromosome removal, so the 4.3-fold figure is the one realized in practice.

Against external Salmon^28^ at production scale, the integrated TranscriptVB quantification agrees at the gene level used by downstream MorPhiC differential-expression workflows: gene Spearman 0.9992 and Pearson 1.0000 on the read counts each tool assigns to every gene. At the transcript level, read-count Spearman is 0.985 and Pearson 0.99998 over all transcripts (transcripts-per-million Pearson 0.9997). These small transcript-level differences localize to per-read processing order under multithreading rather than to biological signal. These and the other modules’ residual discrepancies are traced to their sources and grouped into documented variance classes (Supplementary Table S6), assessed in Section 2.8.

### 2.3 scRNA-seq: parity with Cell Ranger 9 for production adoption

STARsolo’s documented Cell Ranger compatibility stops at Cell Ranger 5. Cell Ranger 7 made intron-inclusive counting the default. Although STARsolo can count intronic reads, its multimapper handling was never revised for the cases that intronic counting creates—small genes and non-coding RNAs that lie within the introns of other genes, where a read’s assignment depends on how a multimapper is resolved. Parameter tuning can improve the concordance, and the Sanger Cellular Genetics group’s STARsolo recommendations (CellGENI ^29^) are the established configuration for approximating Cell Ranger with upstream STAR. But flags can only select the multimapper and overlap rules that STARsolo already has; they cannot add a rule where none exists. The residual disagreements between STARsolo and Cell Ranger arise from artifacts and limitations that became consequential as counting moved beyond exonic reads to include intronic and long non-coding RNA signal.

Poly-G artifacts from two-color NovaSeq and NextSeq chemistry were negligible for exon-only counting but now inflate some gene counts substantially. STAR-core detects and removes them during alignment (`--clip3pPolyG`). A multimapper policy matched to Cell Ranger’s outputs resolves the ambiguous reads common in repetitive intronic and lncRNA regions.

On the public 10x PBMC 10K dataset (Table 1), with both tools run in one session on the same machine, STAR Suite gave barcode Jaccard 0.995, cell Pearson 0.99999, mean per-cell Pearson 0.992, gene Spearman 0.990 and gene Pearson 0.9998 against Cell Ranger 9.0.1, in 16 min 18 s against 63 min (3.9-fold; Tables 2 and 3). The same data run through upstream STAR 2.7.11b with the CellGENI parameters gave gene Spearman 0.950, gene Pearson 0.993 and barcode Jaccard 0.994 (Supplementary Table S1); the difference lies in the gene-level counts rather than in cell calling or per-cell totals (cell Pearson 0.99996 against 0.99999).

The gap is not particular to PBMC. On two further public datasets, a low-coverage glioma line (GSE261042) and a GEM-X 3’ v4 organoid (GSE316010), STAR Suite gave gene Spearman 0.992 and 0.993 and the CellGENI configuration 0.957 and 0.960, at similar Jaccards, and STAR Suite ran 1.5- to 1.9-fold faster than the CellGENI configuration on all three datasets (Supplementary Table S1).

The same genes account for the difference in every dataset: the CellGENI counts over-count LINC00486 16- to 106-fold and under-count highly expressed genes with many pseudogene copies, such as RPL39, H3-3A and SUMO2, to 8–27% of Cell Ranger’s counts, which a matrix of uniquely mapped reads loses; these are the counts that the poly-G trimming (LINC00486) and the multimapper policy (the pseudogene-rich genes) above bring into agreement.

The major remaining disagreement between STAR Suite and Cell Ranger is on cell calls, where Monte Carlo sampling in EmptyDrops ^30^ makes small UMI differences decisive (Section 2.8). On all three datasets the disagreement is one-sided: STAR Suite called 11,863 cells against Cell Ranger’s 11,806 on PBMC, 6,441 against 6,427 on the glioma line and 10,352 against 10,060 on the organoid, and in each case almost every Cell Ranger call lies inside STAR Suite’s set (Supplementary Table S1). The difference is a few extra marginal cells, which downstream filtering removes, rather than calls that STAR Suite missed.

STAR-core also supports 10x’s newer GEM-X OCM (on-chip multiplexing) single-cell chemistry through Cell Ranger-compatible composite barcode handling (`--ocmMultiEnable`), so the same scRNA-seq routines process classic 10x and GEM-X OCM inputs without external preprocessing. RNA-velocity support was a specific request from consortium analysts. STARsolo already writes the raw and filtered spliced/unspliced matrices that velocity tools ^31^ require—output Cell Ranger does not provide, leaving its users to run a separate splice-aware quantification pass. Our contribution is not added capability but added consistency: the STAR-core improvements above— poly-G removal, the Cell Ranger-matched multimapper policy, and intronic counting—now flow through to the Velocyto-compatible matrices under `outs/`, so the velocity layers are built from the same clipped reads, resolved alignments and corrected barcodes and UMIs as the gene-level matrix. The original STARsolo velocity behavior remains available unchanged, as does all prior STAR functionality. The single-cell benchmarks compare against Cell Ranger, which neither removes the Y chromosome nor writes velocity matrices, so both steps are left out of those runs (Methods).

### 2.4 STAR-Perturb: open-source feature processing at production scale

The open-source options for feature barcodes—R or Python scripts, assay-specific implementations, aligner-based workarounds assembled for one experiment at a time—do not scale to production, which left Cell Ranger as the only practical route for CRISPR guide capture in Perturb-seq^2^ and for related assays such as clonal tracking ^32^ and epitope measurement (CITE-seq^33^). We needed an open-source tool that could process feature barcodes at consortium scale—on the MSK 30-KO dataset, for example, matching reads against a LARRY lineage library of 245,979 barcodes alongside 30 CRISPR guides—and that supported the feature-barcode calling methods used in the consortium.

We built the standalone feature-assignment engine `process_features`^34^ to meet this need. It uses a tiered matching strategy: hash tables resolve exact matches and matches within one or two mismatches against the listed feature library, which covers nearly every read; where a library needs a wider search than those tables can hold, fast-Hamming, the exact vectorized Hamming-distance search introduced with STAR-Flex below, takes over (Fig. 3b; Methods). `process_features` is available standalone for users who need feature processing without genomic alignment.

**Figure 3:**
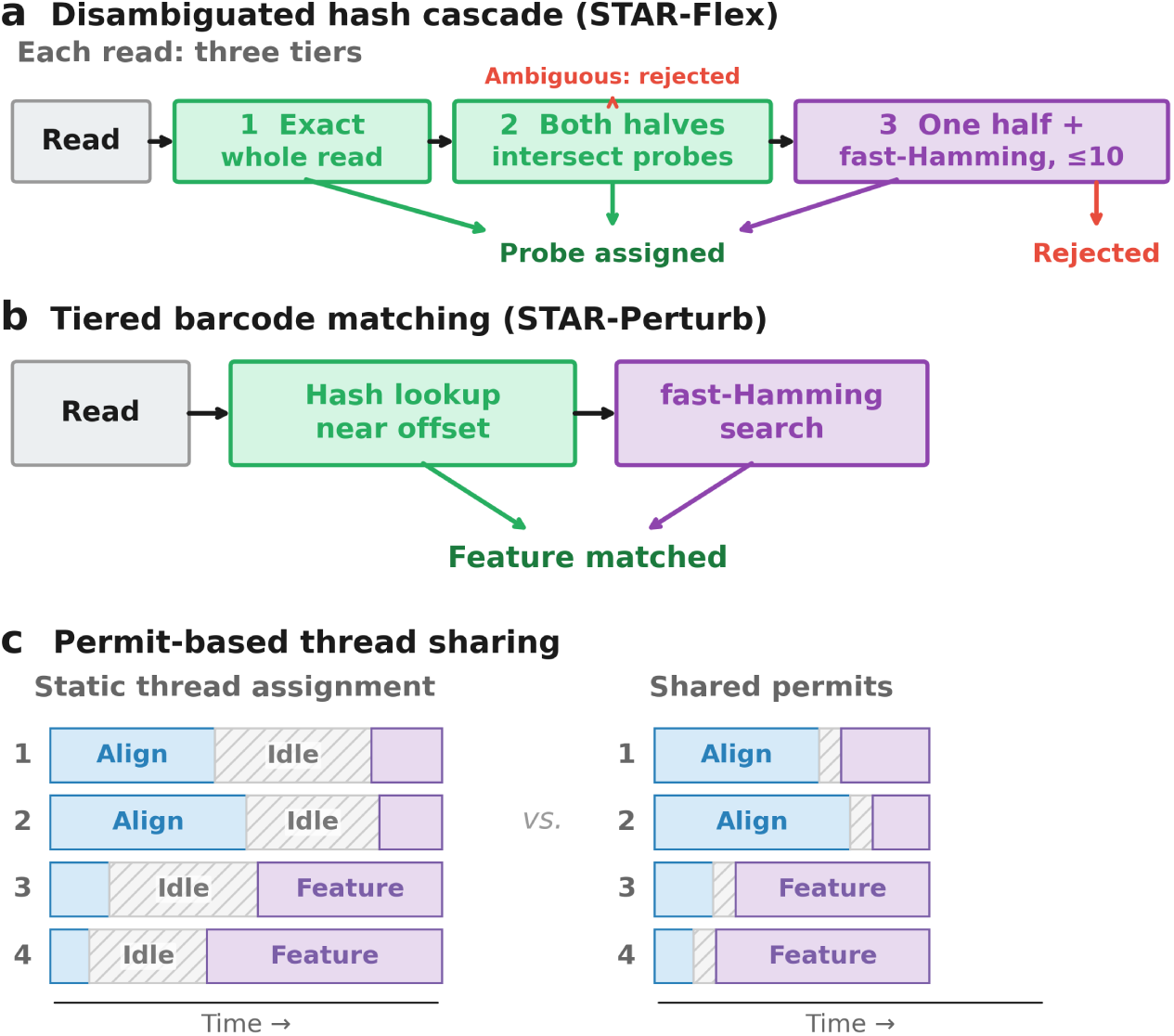
Algorithmic strategies enabled by single-process integration. Panels a and b apply the same matching principles. Reads are matched by hash lookup wherever possible, because a lookup is fast. Tables are kept as small as the task allows, for memory locality. Where a hash cannot be used or would be inefficient, at larger Hamming distances or for longer sequences, fast-Hamming, a vectorized exact Hamming-distance search, finds the best match. **a,** STAR-Flex probe assignment, the disambiguated hash cascade, as each read passes through it. The tables it queries—one of the exact probes, and one per 25-base half holding every sequence within one mismatch of that half, each entry recording the probes that can produce it—depend only on the probe set and, like a genome index, are built once, outside the timed runs (Methods). A read is assigned by (1) exact lookup; (2) lookup of both halves, when their probes intersect in one probe; or (3) when only one half identifies a single probe, fast-Hamming scoring of the other half, accepting at most ten mismatches over both halves (Cell Ranger’s documented tolerance ^35^). Reads whose halves disagree, or that match an ambiguous sequence, are rejected. **b,** STAR-Perturb feature-barcode matching. Tables of each listed feature and its variants within one and two mismatches, near the expected offset, assign most reads; validation marks variants that more than one feature could produce as ambiguous; fast-Hamming handles larger distances and longer sequences, where variant tables would be too large.

Running `process_features` and the STAR aligner as separate processes immediately exposed a second production pain point: both have characteristic lag periods—disk I/O stalls during sequencing-file decompression and barcode-table loading—during which the other has work it could be doing. Integrating feature processing directly into the STAR executable removed this idle time. Inside a single process, threads are reassigned dynamically between alignment and feature **c,** Permit-based thread sharing, the general mechanism by which the stages of one process share its threads. With a static assignment (left), each thread is tied to one workload and idles (grey) when that workload stalls. With shared permits (right), work runs while it holds a permit, so threads pass to whichever stage has work waiting, here from alignment to feature assignment, so the idle stretches shrink and the run finishes sooner. In STAR-Flex the same permits are used for block decompression and cell calling (Methods). Schematic, not measured. assignment as the run progresses (Fig. 3c): a permit-based thread scheduler requires each unit of alignment or feature work to hold one of a fixed pool of permits, so when either workload stalls its permits pass to the other and the available cores shift with them (Methods). This function-call-level interleaving is much finer-grained load balancing than is achievable across pipeline stages coordinated through Nextflow, Kubernetes, or the Martian pipeline framework that Cell Ranger documents as its scheduler ^36^.

Across the A375 and MSK 30-KO benchmarks (Table 1), STAR-Perturb completed in 2 min 26 s and 27 min 6 s respectively, corresponding to 4.7- and 6.2-fold speedups over Cell Ranger 9.0.1 (Table 2; Supplementary Table S2). The largest gain was on the MSK dataset, where STAR-Perturb processed gene expression, CRISPR guide capture, and the LARRY lineage library simultaneously in a single run while Cell Ranger required separate runs for each feature barcode set. Simultaneous processing of expression, guide, and lineage libraries in one pass is itself a capability that no other tool, open or proprietary, currently provides.

Production data centers were already using Cell Ranger’s Gaussian mixture model methodology for Perturb-seq, so STAR-Perturb mirrors the GMM-based feature calling for CRISPR guides. For lineage barcodes we added a dominance rule, better suited to high-multiplicity lineage barcoding, where the relevant question is whether one LARRY barcode dominates a cell’s lineage signal. This mirrors what the data production centers used previously, and integrated into the run it reproduces the production calls exactly on the MSK benchmark (Methods). On A375, CRISPR calling agreed exactly on all 1,076 cells carrying a guide call in both outputs; on the larger MSK dataset exact CRISPR call agreement was 99.0% with per-guide feature-UMI Pearson correlations of 0.999 (Table 3; Supplementary Table S2). Where STAR-Perturb and Cell Ranger diverge, part of the difference is valid feature matches that STAR-Perturb detects and Cell Ranger’s feature-count tables omit: our match tables and direct inspection of the original FASTQ files confirm the reads are present and match the listed feature sequences exactly. The remaining differences—connected-component UMI correction ^37^ (versus the single-base UMI correction Cell Ranger documents^38^) and Monte Carlo variation in EmptyDrops ^30^ cell-calling—are localized in Table 5 (Section 2.8).

### 2.5 STAR-Flex: a disambiguated hash cascade for Flex processing without alignment

10x Genomics Fixed RNA Profiling (Flex) is a probe-based, lower-cost single-cell RNA-seq assay that works on fixed and formalin-fixed, paraffin-embedded (FFPE) material and multiplexes several samples in one lane. Each read carries a 50-base probe sequence and, on the same mate, an 8-base sample tag from RNA-templated ligation (RTL), so probe assignment, sample-tag demultiplexing and cell calling are tightly linked. In a multiplexed dataset a cell is a barcode together with a tag, with several tags often belonging to one biological sample. What makes probe assignment hard is the tolerance the assay requires. A read may carry up to ten mismatches across its two 25-base halves, far more than an enumerated-variant table can hold: a single 25-base half has 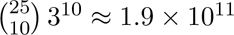 variants with exactly ten mismatches, a whole probe has 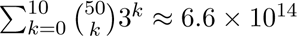 variants within ten, about 3.5 *×* 10^19^ across the more than 50,000 probes of a probe set, against 25 *×* 3 = 75 single-mismatch variants per half. Some per-read scoring beyond lookup is therefore unavoidable; Cell Ranger’s documented probe aligner supplies it by alignment^35^, and the one open-source processor, the recent standalone k-mer tool cyto ^39^, by k-mer matching.

We arrived at STAR-Flex’s method in three steps, each recorded in the repository’s commits and design documents. We began with alignment, leveraging the integration with the STAR aligner. Reads were aligned against a hybrid reference carrying a pseudo-chromosome for every probe. That matched Cell Ranger’s results very well but required separate passes and large intermediate files. For speed we moved to an alignment-validated hash: every single-mismatch variant of every probe was enumerated once, while the cache was built, aligned against that reference, and kept only if it mapped uniquely to its probe and there was no better alignment from genomic sequences. Establishing that cell-barcode and UMI resolution does not depend on the sample tag let tag detection run inline in the same pass over the reads (Methods), and this gave the first single-process pipeline. On the deeply sequenced JAX SC2300771 dataset it agreed closely with Cell Ranger, and it was already far faster: in release 1.8.0’s benchmark record the alignment-validated route processed JAX SC2300771 in 10 min 18 s and full alignment in 23 min 22 s, against 59 min for a contemporaneous Cell Ranger 9 run with secondary analysis enabled (5.7- and 2.5-fold).

Testing at lower coverage on multiplexed samples showed that this was not enough. On the public 10x 320k dataset, sequenced more shallowly and with two tags per sample, the cache’s conservative rejection of variants that also resemble genomic sequence shifted near-threshold cells into and out of the called set. Rerun with the current cell caller, the alignment-validated cache agrees with Cell Ranger at a pooled Jaccard of 0.990 on JAX but 0.933 on 320k, against 0.988 and 0.974 for the half-probe rule described next (Supplementary Note 3, Table S10).

At this point we decided to implement the procedure 10x documents for its probe aligner ^35^ directly, as a disambiguated hash cascade (Fig. 3a). We kept our exact 50-base hash for exact matches but used two 25-base half-sequence caches. For each probe we generated all single-substitution variants of the 5*^′^* 25 bases, recording with each variant the probes that can produce it, and repeated the enumeration on the 3*^′^* half. The check that alignment used to perform now happens when a read is looked up rather than when the cache is built: a read whose two halves have exactly one probe in common is assigned to it, and one whose halves have more than one is rejected as ambiguous. If only one half can be matched, we determine the exact Hamming distance to the other half of the probe. The read is accepted when the two halves together carry at most ten mismatches. This matches the 10x documented procedure, except that where 10x scores the second half by a bounded edit-distance alignment, we use the Hamming distance.

For the Hamming distance determination we use fast-Hamming, our vectorized exact Hamming-distance search, which returns the exact distance between two packed sequences in a few instructions (Fig. 3a; Methods). It is also used in feature-barcode assignment, where it is again the fallback behind the hash tiers. With our current methodology, no read is aligned and no genome is loaded, greatly reducing the memory footprint (Scratch storage and Flex on spinning disk, below). The original alignment-validated cache and full alignment remain available behind a compatibility flag (--flexLegacy yes) for users who are concerned about genomic cross-assignments (Discussion).

For Flex we ran additional benchmarks that vary the input format and the storage medium. Because cyto is designed around BINSEQ (CBQ) input, Table 2 carries CBQ rows for Flex alongside rows for the identical original files, so that the two tools are compared on the input each prefers as well as on the same files (Methods). Storage is an important variable because Cell Ranger’s Flex run on the 320k dataset needs terabytes of dedicated fast scratch (below). Our server does not have that much SSD, and many laboratories do not either: it is why the 320k comparison had to be run on a cloud instance, and why sites that process Flex at this scale keep their working files on spinning disk or network storage. The three Flex benchmarks were therefore repeated with every file on spinning disk, so that the comparison also reflects the storage on which Cell Ranger will actually be run in many environments. The input routes are described with STAR-core (Section 2.1). The storage measurements follow (Supplementary Note 2, Table S9).

STAR-Flex creates the same per-sample filtered matrices produced by Cell Ranger 9.0.1 after cell calling. On the two single-tag datasets it called slightly more cells than Cell Ranger and missed almost none of Cell Ranger’s—4 of 20,419 on JAX SC2300771 and 5 of 38,444 on GSE325982—for a pooled barcode Jaccard of 0.988 and 0.994, and 0.970 to 0.997 sample by sample. The counts on the shared cells were very close: mean per-cell Pearson 0.995 to 0.999, cell Pearson above 0.9999, gene Spearman above 0.9998 and gene Pearson above 0.999998 in every sample (Table 3; per sample, Supplementary Table S3). Both datasets ran about 17-fold faster than Cell Ranger 9.0.1 from the original FASTQ—3 min 20 s against 57 min 6 s on JAX, 4 min 17 s against 74 min 6 s on GSE325982—and 20- and 41-fold faster from CBQ, in 24 and 18 GiB of memory (Table 2).

The public 320k dataset comes from the newer GEM-X chip generation, which repartitions many more cells per lane—320,000 cells across 16 tags. STAR-Flex processes both chip generations with an identical configuration. This dataset has 7.3 billion read pairs across eight tissues, two tags to a tissue (Table 1). Because of the amount of space Cell Ranger needs for its intermediate files, we ran this benchmark on an AWS m6id.16xlarge (Intel Xeon Platinum 8375C, 32 virtual CPUs, 128 GiB of memory) with two local NVMe SSDs in RAID0, an instance of the lab server’s size (Methods). With that configuration STAR-Flex completes in 17 min 36 s at 32 threads with 72 GiB peak memory, 22.6-fold faster than Cell Ranger 9.0.1 (6 h 38 min), with called-cell Jaccard 0.974 against the Cell Ranger 9 reference (333,411 STAR cells against 325,410), 0.958 to 0.984 across the eight samples, and mean per-cell Pearson 0.9990 to 0.9992.

We measured cyto on all three benchmarks on the same hardware as STAR-Flex (Supplementary Note 1, Table S7). The two tools handle a multiplexed sample differently: STAR-Flex, like Cell Ranger, calls cells per biological sample, pooling a sample’s tags into one model (Methods), while cyto, following its documented route, filters each tag separately and concatenates the per-tag calls. On the single-tag JAX SC2300771 and GSE325982 samples this makes no difference and the two are close, at Jaccard 0.985 against 0.988 and 0.9933 against 0.9939. On the 320k samples, each spread over two tags, cyto’s called-cell agreement with Cell Ranger falls to 0.696–0.952 per sample (pooled 0.883) against 0.958–0.984 (pooled 0.974) for STAR-Flex, while the counts agree closely for both: on the 314,388 cells all three tools call, so that the two are measured on identical cells, the mean per-cell Pearson with Cell Ranger is 0.9991 for STAR-Flex and 0.9958 for cyto (Supplementary Table S8). STAR-Flex was also 1.1- to 1.4-fold faster on every leg, from the original files and from binary input alike, and used less memory (Supplementary Table S7; Scratch storage and Flex on spinning disk, below). We ran cyto in h5ad mode throughout because only this mode performed cell calling (Supplementary Note 1).

### 2.6 Scratch storage and Flex on spinning disk

STAR-Flex runs on a modest server without trading disk for memory: it writes almost nothing beyond its outputs, and it does so in less memory than cyto and within the memory Cell Ranger was given. The difference in disk is due to Cell Ranger’s splitting into chunks that are written to disk to be processed in parallel. These intermediates are removed only once the stages that depend on them have completed ^36^. On the 320k run the alignment stage alone ran as 100 chunks. That trades parallelism for disk usage. In contrast, STAR-Flex parallelizes with threads inside one process and keeps its working state in memory.

The difference is large. On the 320k dataset Cell Ranger’s intermediates peaked at 1,805 GiB, cyto at 218 GiB and STAR-Flex at 16.9 GiB, essentially its retained output; the ordering is the same on the two smaller datasets, where STAR-Flex peaks at 2.5 to 4.6 GiB (Supplementary Table S9). Disk use is reported as this peak, the most a run holds at once, because that is what a site has to provide. Memory follows the same pattern: STAR-Flex peaked at 18 to 72 GiB across the three datasets against 37 to 115 GiB for cyto, with cyto under a 120 GiB limit and Cell Ranger allocated 120 GB (Supplementary Table S7). For scale, 10x’s system requirements for Cell Ranger state a minimum of 64 GB of memory, with 128 GB recommended and more for larger datasets, and 1 TB of free disk ^40^: STAR-Flex’s 72 GiB on the largest public Flex dataset sits inside that memory range, while its disk use is two orders of magnitude below the disk minimum. For datasets larger than memory allows its cell-barcode buckets can spill to disk, and none of these benchmarks needed it. Repeating the three Flex benchmarks with every file on spinning disk, on a cloud instance matched to the NVMe instance in size and processor throughput, leaves the ordering of the three tools unchanged (Supplementary Note 2; Supplementary Table S9): STAR-Flex remains 12- to 20-fold faster than Cell Ranger from the original files, and its margin over cyto widens from 1.1– 1.4-fold on NVMe to 1.5–3.9-fold on spinning disk, consistent with the disk each tool needs for the same work. These comparisons hold the storage the same for every tool, which is the conservative choice: on sets large enough that Cell Ranger’s scratch no longer fits on a site’s SSD, the realistic placement is STAR-Flex on local NVMe against Cell Ranger on an array (Discussion).

### 2.7 STAR-SLAM: integrated NTR estimation for production SLAM-seq analysis

SLAM-seq^5^ measures RNA turnover by metabolically labeling newly transcribed RNA with 4-thiouridine (s4U), which introduces T*>*C conversions during reverse transcription. The ratio of converted to total reads at each gene—the new-to-total RNA ratio—provides a direct readout of transcriptional dynamics and mRNA half-life, complementing steady-state expression measurements. SLAM-seq was among the assays generated by MorPhiC data production centers. Production first used SLAMDUNK^41^, but it relies on its own aligner, does not handle the paired-end reads MorPhiC generates, and estimates NTR with a less robust statistical model. Because the consortium needed differential-expression analysis alongside turnover, we moved to GRAND-SLAM^42^ (built on the GEDI genomics platform), which uses STAR for alignment and applies a stronger statistical model for conversion detection and NTR estimation^43^. That workflow—aligning with STAR, then handing the alignments to GRAND-SLAM—nonetheless carried three production problems we needed to address.

First, *logic drift*. Because GRAND-SLAM operates on STAR’s output files rather than within the aligner, it re-derives gene assignments and counting decisions independently of how STAR made them; as the aligner evolves between releases, the downstream tool’s approximations diverge silently from the alignment-level decisions, which can introduce reproducibility differences between expression counts and NTR estimates. Second, *paired-end counting for downstream DESeq2*. MorPhiC’s SLAM-seq data is paired-end, and standard DESeq2^24^ differential-expression workflows expect the same paired-end-aware counts to be used for both the differential expression call and the NTR-based turnover call. GRAND-SLAM does not produce that paired-end-aware count table natively, so it required additional manual steps. Third, *true variant masks across multi-sample experiments*. For well-characterized cell lines such as KOLF2.2J we have high-confidence variant calls and want to mask known SNPs using those calls rather than estimating SNP-site likelihoods from the SLAM-seq data itself, but applying per-sample VCF masks is awkward in GRAND-SLAM when handling the many samples of a production SLAM-seq experiment, and required additional scripting.

STAR-SLAM addresses all three by performing T*>*C mutation detection, SNP masking, and NTR estimation directly within the aligner’s counting loop. The same gene-assignment and overlap-resolution logic governs both expression quantification and conversion calling, eliminating the logic drift between expression counts and NTR values. Paired-end handling is consistent with how the aligner counts reads for downstream differential expression, so the SLAM-seq count tables and the standard expression count tables come from a single pass and feed into DESeq2 without reconciling NTR values with the paired-end gene counts. External variant masks are applied per sample as ordinary inputs, with no scripting around the tool. STAR-SLAM also adds variance-based auto-trimming, a data-driven alternative to the manual or default trimming windows other tools require. The implementation—the binomial-EM NTR model, the auto-trimming algorithm, the internal SNP-detection fallback, and a GEDI-compatibility mode for controlled migration—is detailed in Methods.

On the GRAND-SLAM benchmark (Table 1), STAR-SLAM achieves NTR Pearson of 0.994 to 0.999 across read-count thresholds (Table 3; Supplementary Table S4), and its output follows the GRAND-SLAM column schema so existing downstream tools run unchanged.

We do not report a speed comparison for SLAM-seq, for two reasons. GRAND-SLAM relies on an aligner such as STAR for its alignments but has its own count generator in GEDI, so the time of the pipeline it belongs to is set largely by the aligner and the steps around it rather than by GRAND-SLAM itself. And the two methods handle known variants differently: STAR-SLAM masks the variants listed in a VCF for each sample, whereas GRAND-SLAM calls variant sites itself from the SLAM-seq reads, so a GRAND-SLAM pipeline that uses the same variant calls needs extra steps that are not part of GRAND-SLAM. With this much of the work done differently, a timing would not compare like with like.

### 2.8 Clean-room reimplementation, and the residual differences that remain

Cell Ranger’s source is public, but its license bars using it to build a derivative, so every Cell Ranger-compatible component was reimplemented clean-room. We used the descriptions from 10x’s published documentation when available. Gaps in the methodology were inferred from the observed outputs of matched STAR Suite and Cell Ranger runs. For the open-source modules (Cutadapt, samtools, Salmon and GRAND-SLAM) we copied no code either, but we were free to read it, and did so to resolve gaps in the documentation.

Adapter trimming reproduces its reference byte for byte. Coordinate sorting reproduces every SAM record, although the BGZF block boundaries of the compressed file are not identical across builds. For most of the others the outputs are very similar but not identical, and Table 5 maps each module’s residuals to the variance classes catalogued in Supplementary Table S6, indexed in brackets there.

The first source of residuals is intentional methodological difference from the reference. An example is Flex probe assignment. Where a read anchors on one probe half, 10x’s documented aligner scores the other half by a bounded edit-distance alignment^35^; we score it by the exact Hamming distance. We prefer the Hamming distance because the assay’s ligation design makes substitutions the dominant error, and because the distance is easily computed using fast-Hamming. A read carrying an indel in that half is therefore accepted by Cell Ranger within its ten-edit budget and rejected by STAR-Flex [12], and the close agreement with Cell Ranger on all three Flex datasets shows how rarely that happens (STAR-Flex).

A second related source of residuals occurs when the clean-room implementation is unable to infer all of the underlying rules. For example, we do not know the exact policy that Cell Ranger uses to resolve multimappers in scRNA-seq since we cannot look at the code. Instead, we infer the rules that we can and fill in any gaps with principled rules such as preferring exonic matches to intronic matches. In both these cases, the residuals are due to principled and recorded differences in policies and are not errors per se.

The third contributor to residuals is steps that are stochastic or depend on the order in which reads are processed. An example is EmptyDrops-based cell calling, used by STAR Suite and Cell Ranger, which relies on Monte Carlo sampling of the ambient profile [1]. In bulk RNA-seq, Salmon fixes its library type from a running sample of the first reads it sees [4], while TranscriptVB’s counts depend on the order in which its threads deliver reads [5], so the two agree exactly only when the run is single-threaded and the order is matched. These factors produce small random differences with minimal biological effect, but they prevent full parity even when everything else is matched.

### 2.9 AI-assisted development built a maintainable platform

STAR Suite adds 183,606 lines of C/C++ to the 28,228-line STAR codebase (counted by `tools/count_cpp_lines.sh`; Methods) and, apart from OpenSSL’s libcrypto, no dependencies beyond what upstream STAR already uses. A single developer-scientist built it through the “Human-Architect, AI-Implementer” cycle of Fig. 4 (Methods). Each feature began as an architectural plan, written by the developer with an AI agent. The plan fixed the module boundaries and file ownership, the data structures and their memory behavior, and the points at which new code meets the existing aligner. It also established the parity metrics and outputs the result would be judged against. Wherever production data covered the feature, the developer supplied gold standards built from it. Agents then implemented the plan, wrote and ran unit tests and the end-to-end regressions against those gold standards, and the developer reviewed and redirected the work until the parity metrics passed. Table 6 records, for each component, what the developer contributed and what the agents did. All of the new code was written by AI except `process_features`, whose roughly 8,700 lines predate the project and used AI only for boilerplate and code completion.

**Figure 4:**
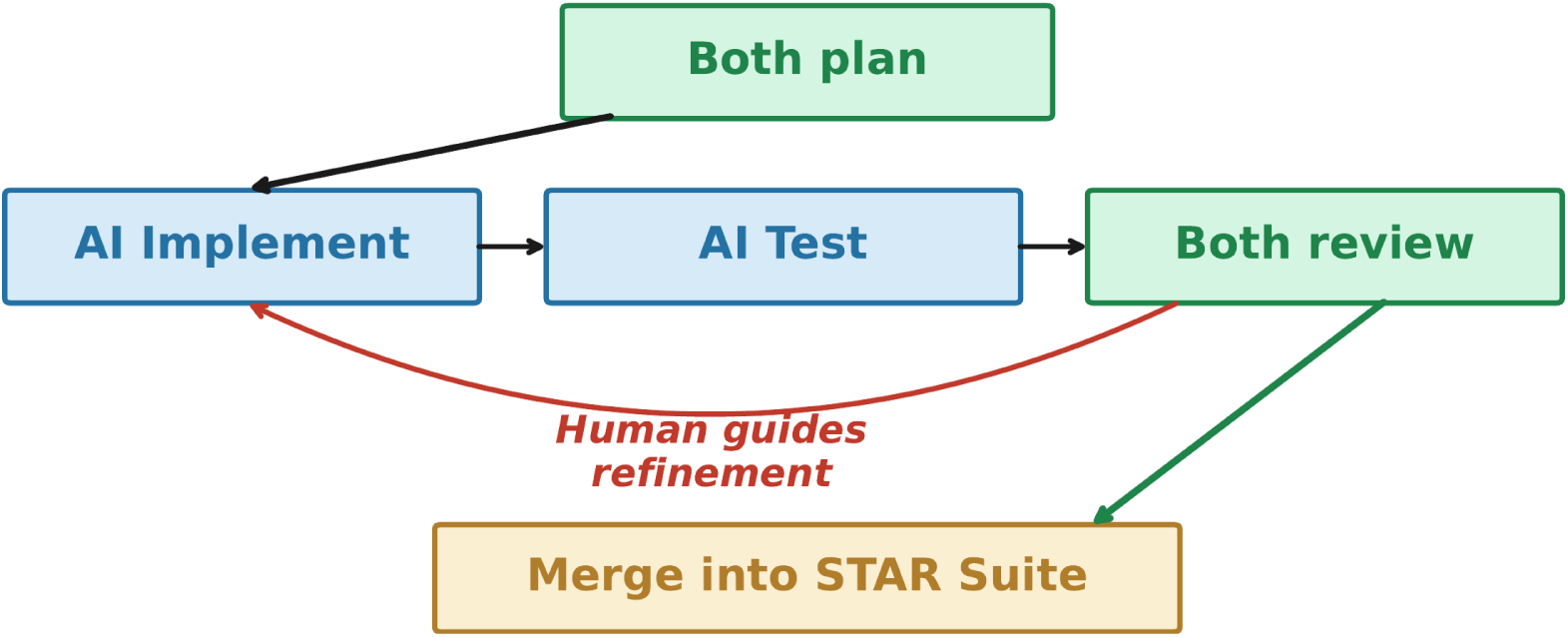
Human-Architect, AI-Implementer development workflow. Color marks who does the work: blue, AI; green, both; gold, a STAR Suite artifact. The red return arrow is the human-guided step. A review that passes is merged into STAR Suite. A review that fails returns for another round of AI refinement, guided by the human.

**Table 6.** Division of labor between human and AI in STAR Suite’s development. The core algorithms, integration targets, overall architecture, and end-to-end tests were the human’s; AI agents produced most of the integration code, the synthetic tests, the interface contracts, and the parameter sweeps, and co-designed the permit-based thread scheduler, the one component whose implementation relied on AI. For the other components a viable implementation existed or would have existed without AI. There, AI chiefly made the work faster to build and more robust.

| Capability | Human contribution | AI contribution |
| --- | --- | --- |
| fast-Hamming | Novel algorithm and standalone implementation (`process_features`, largely predating AI) | Integration into STAR |
| Flex probe assignment (disambiguated hash cascade) | Design of the methods: first the hybrid pseudo-chromosome reference and the alignment-validated cache, then the disambiguated cascade and the decision to adopt the documented two-half tolerance | Implementation of the alignment method, both caches and the grouped cell caller; parameter sweeps |
| Permit-based thread scheduler | Concept and aim | First working implementation; extensive co-design |
| Adapter trimming | Target: Cutadapt-equivalent behavior | In-process C/C++ implementation; parity tests |
| Y-chromosome removal | Target: inline removal during alignment | Inline BAM/FASTQ splitting; tests |
| SLAM-seq auto-trimming | Target: variance-based auto-trimming | Implementation; tests |
| VCF-based background removal | Target: SNP masking from external calls | Masking implementation; tests |
| Codebase integration | Architecture, specifications, end-to-end tests | Most integration code; synthetic tests; interface contracts |

Figure 5 shows a codebase that has been maintained and updated in production for eight months from January 2026, when the feature-barcode and Flex work, begun in standalone repositories in 2025 (process features has been public since December 2024), was brought together in the STAR-suite repository. It was built out rapidly in early 2026 and has since been optimized, maintained and integrated with companion multiomic tooling across 25 public releases.

**Figure 5:**
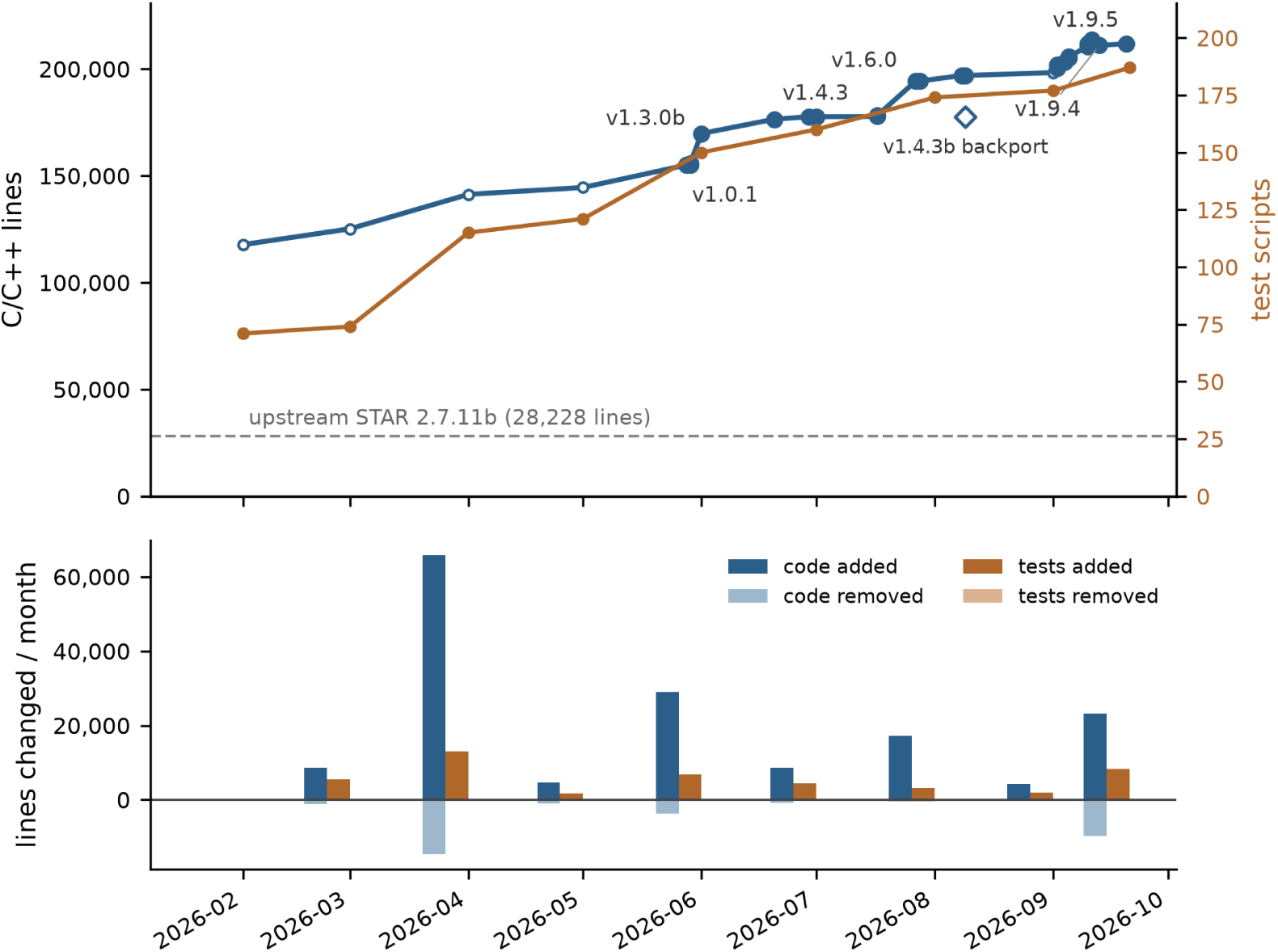
A maintained, evolving codebase. The record begins when the standalone feature-barcode and Flex repositories were brought together in January 2026, and the series is plotted from the following month start. **Top:** total C/C++ lines (counted by `tools/count_cpp_lines.sh`, vendored libraries excluded) at each of the 25 public releases, against the 28,228-line upstream STAR 2.7.11b base (dashed). Open circles are month starts before and between releases, and the open diamond is v1.4.3b, a fix backported to the 1.4 line for users of that release. The near-flat spans are surgical maintenance releases: the three releases from the labelled v1.4.3 through v1.5.0 added 637 lines between them. The later steps, such as v1.6.0, are maturity-phase work made into the established codebase: performance optimization, a major bug fix, and integration with companion multiomic tooling and HPC-scale deployment. The September 2026 releases (v1.8.0–v1.9.5) carry the parallel BGZF reader, the half-probe Flex route, the Flex memory rework and the integrated lineage caller, and v1.9.4 removed more code than it added. The second axis tracks the regression-test scripts, growing from 71 to 187 over the same period. **Bottom:** lines added and removed per month, for code and for tests, with September 2026 shown to the 20th: roughly 32,000 code lines and 1,410 test lines were deleted over the project—the codebase and its tests are continuously reworked, not appended to.

Before a release is published it passes a battery of tests: continuous integration on every change, validation of the built binaries and packages in the release workflow, and fixture and production-scale comparisons run before the tag is cut (Methods). Each release’s notes state what it fixed, changed and validated. Releases can also retire and clean up superseded legacy code—roughly 32,000 lines of code and 1,410 lines of tests have been deleted over the project.

The current repository holds 814 commits, 187 regression-test scripts, release notes for each release, and 311 benchmark records. There are 275 working documents—the architectural plans, runbooks and handoff notes that guided AI agents. The repository is unsanitized—one can track how the codebase has evolved, including bugs and dead ends. This is important for AI-assisted development: it prevents agents from repeating mistakes and re-attempting failed approaches, and the complete record gives new agents the context needed for further modification of the codebase and facilitates the resolution of bugs.

### 2.10 One production interface for humans and agents

The result is a single self-contained binary that human operators and AI agents both work with from one shared set of typed recipes, which agents reach through a Model Context Protocol (MCP) server ^44^ and people through the browser-based Launchpad, so the two routes render the same validated command and cannot drift apart (Fig. 6). In production, the scripts that run it are kept in morphic-provenance together with the exact commands, inputs, outputs and deliveries of each run, and a later correction is added with its reason rather than written over. The interface and the two layers of recipes and records are detailed in Methods.

**Figure 6:**
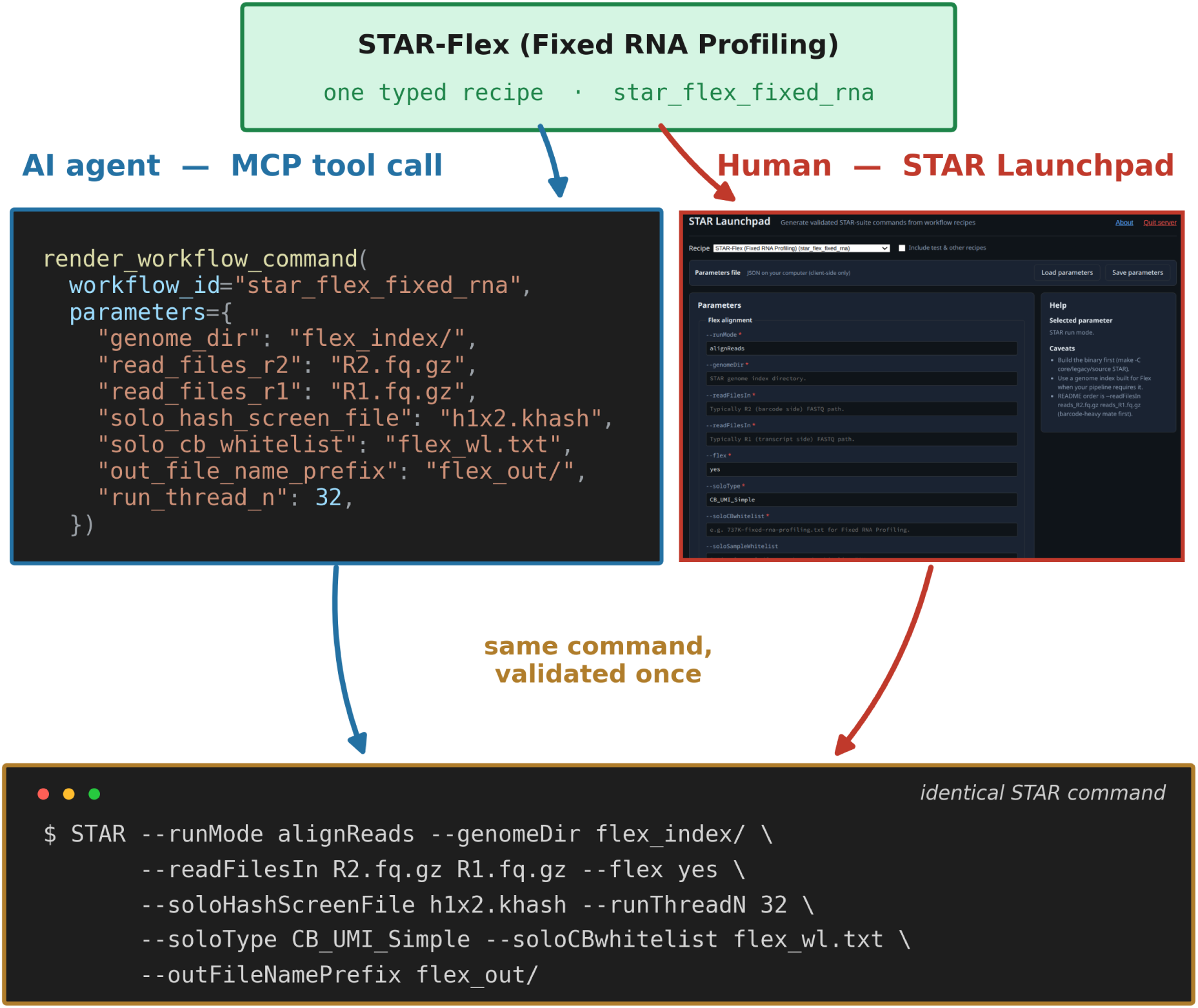
One recipe, two front doors, one command. The STAR MCP server holds one typed workflow recipe—the parameter contract—and presents it two ways: to an AI agent as a schema-validated tool call (left), and to a human operator as the same recipe rendered into a browser-based Launchpad form (right; screenshot from the STAR-suite documentation). Shown here for the STAR-Flex recipe (star flex fixed rna). Either route returns the filled parameters and renders the identical validated STAR command (bottom). The same mechanism is used for every recipe, and recipes compose into multi-assay pipelines.

## 3 Discussion

The several-fold gains for bulk, scRNA-seq and Perturb-seq and tens-fold gains for Flex in Table 2 are not from simple optimizations, from trading accuracy for speed, or from translation to compiled C++. They come from new techniques that process the data more efficiently, and from integration into STAR, which removes the overhead of interprocess orchestration and lets the stages coordinate internally. The disambiguated hash cascade and fast-Hamming in effect assign a Flex probe through table lookups and a single vectorized comparison. The tiered hash search in process features, with fast-Hamming for larger distances, does the same for feature barcodes.

In our original bulk RNA-seq pipeline, Cutadapt (through Trim Galore) decompressed the input FASTQ files and recompressed them for the aligner to decompress again. Y-chromosome removal decompressed and recompressed FASTQ and BAM files, and Salmon operated on BAM files. All of these intermediate steps are avoided by integrating the functions and operating on the internal data structures. There is also no need to manage, load and unload container images to keep the dependencies reproducible. Integration also allows feature-barcode assignment, alignment, cell-barcode mapping, transcript counting and cell calling to be coordinated internally, so that threads can be assigned dynamically to the procedure that can best use them. In SLAM-seq the gain is correctness rather than speed: conversion calling shares the aligner’s own gene assignment, so the turnover estimates and the expression counts a differential-expression workflow uses cannot drift apart, as they do when a downstream tool re-derives them from the alignments (Results). All of these improvements were made without sacrificing accuracy, while preserving every existing function and with essentially no new dependencies.

The order-of-magnitude Flex speedups come with significantly lower disk use, which is an advantage in its own right. STAR-Flex writes little beyond its outputs while using less RAM than cyto and running within the memory 10x recommends for Cell Ranger. On the largest set tested, the 320k set, STAR-Flex’s peak disk use was 16.9 GiB, compared to 218 GiB for cyto. Cell Ranger’s chunked, disk-backed parallelism creates and deletes terabytes of temporary files with a peak disk use of 1,805 GiB (Results). Provisioning multiple terabytes of NVMe SSD per run is a decision that many sites will not or cannot make. On the spinning or network storage they use instead, what a tool writes and re-reads converts directly into elapsed time (Supplementary Note 2). The speedups in Table 2 and Supplementary Table S9 compare the tools on the same storage. In practice STAR-Flex’s working set fits on an SSD while Cell Ranger’s scratch often has to go on spinning disk, and under that placement the 320k run takes 17 min 36 s against 9 h, a 31-fold difference.

We have noted that some of the residual differences (Section 2.8, Table 5) are due to deliberate policy choices. The goal of parity with Cell Ranger outputs is driven by the desire to ensure that users switching over to STAR Suite will obtain the same downstream biological results. However, we did not copy code—ours was a clean-room implementation with legitimate points of disagreement on probe scoring, multimapper handling, feature assignment and, for feature barcodes, UMI correction. Our choices are all documented and produced small deviations. In one case we adopted Cell Ranger’s approach rather than our own: STAR-Flex now uses the half-probe matching strategy that Cell Ranger documents. Previously, we had used a more conservative strategy that checked probe assignments against competing genomic assignments, which gave excellent agreement on high-coverage datasets such as the JAX dataset but much lower agreement on the lower-coverage 320k dataset (Supplementary Note 3, Table S10). It is quite possible that our previous strategy is biologically more accurate. These questions are beyond the scope of this manuscript, but they can now be pursued, because the source code for both approaches is open.

The architecture extends in two directions. Downstream, analysis steps that are conventionally done in Python or R packages can be built into the binary and run on the structures already in memory rather than exchanged through files. Guide calling is the clearest case. STAR-Perturb reproduces Cell Ranger’s Gaussian mixture calling, and what follows it—generalized linear models of the kind SCEPTRE ^45^ uses to test which guides changed expression—is applied by a separate scripting layer that reads the matrices back in. Built into the same process, a caller and the test that consumes it could be refined together and interactively: a threshold changed, the test repeated and the effect seen immediately, which is the kind of repeated pass over one dataset that a scripting intermediate makes slow. The suite also extends across modalities. Our companion preprint describes Chromap Suite, which adds ATAC-seq for multiomic experiments (bioRxiv https://doi.org/10.64898/2026.06.02.729736); it links into the same executable, and the permit-based scheduler already shares one pool of threads across the RNA and ATAC workloads of a joint run (Methods).

STAR Suite is one binary so that people and AI agents can run it in the same way. Biologists, clinicians and data scientists now routinely use AI agents to plan analyses, generate processing scripts and interpret results, and an agent given one command with typed parameters can assemble a valid run without discovering and orchestrating a chain of tools. Because one invocation processes a sample, the script that runs it is compact and readable—a recipe naming its parameters, inputs and the STAR Suite commit—and the record of a run is small enough to be complete. MorPhiC runs this way: STAR Suite is the consortium’s standard transcriptomics processor, its operators use AI agents to write the scripts that run it at scale, and every one of those scripts is kept, versioned, in the public `morphic-provenance` repository^46^. Anyone can read the exact commands and scripts that produced a released output regardless of whether an agent or human created it.

A reengineering of this scale, integrated into a mature codebase by a single developer-scientist, was tractable only through human-directed AI software engineering, disciplined by architectural plans, gold-standard benchmarks from production data and deterministic regression tests (Fig. 4; Table 6). What distinguishes this from the many tools now built with AI assistance is not the speed of construction but the eight months of maintenance in production that followed, with an open public record. The same AI-assisted workflow that built the suite has since fixed its bugs, replaced its methods, cut its memory and retired its legacy routes without changing the outputs its users depend on: the 320k Flex run’s peak fell from 176 GiB before the memory rework to 75 GiB after it, and stands at 72 GiB in the release benchmarked here. This kind of reengineering is now within reach of individual academic groups, not just single-vendor teams. We have already applied the same approach to Chromap Suite, the ATAC-seq processor described above.

That a codebase this large can be maintained with AI assistance is largely due to the nature of bioinformatics data. Bioinformatics tools are often published based on the results of a few datasets because those datasets are rich and varied. A relatively small number of carefully chosen regression tests catches most errors and lets AI coding agents iterate quickly while generating fixes and new features. However, human oversight is needed to guide this process. The problem of maintaining large academic codebases is not new—STAR’s last release, 2.7.11b, was in January 2024, illustrating the limits of the current single-maintainer model. Our hope, and the purpose of the extensive documentation and open code, is that the community, using AI tools, will engage in maintaining and extending STAR Suite.

## 4 Conclusions

STAR Suite integrates the processing of five transcriptomic assays into a single binary. This not only enhances performance by eliminating files written between stages; it also simplifies deployment of processing pipelines by agents and humans and makes analytical workflows easier to audit. The same design extends further: analysis steps now run through Python or R packages, such as the models that test which guides changed expression, can be integrated in the same way, and other modalities added alongside, as our companion ATAC-seq preprint shows. The code is fully open under the MIT license, with the first production-ready open-source Perturb-seq feature-barcode processing and an open end-to-end Flex pipeline. AI agents can freely incorporate STAR Suite into their procedures, and the community can maintain and improve the methodology without navigating a restrictive end-user license.

STAR Suite introduces new methodologies: fast-Hamming, a vectorized exact Hamming-distance search used for Flex probes and feature barcodes; a disambiguated hash cascade that assigns Flex probes without alignment; a permit-based thread scheduler that allocates threads between alignment and feature assignment; and variance-based auto-trimming for SLAM-seq conversion calling. Along with the benefits of integration, these yield significant speedups of 3.9- to 6.2-fold over Cell Ranger for scRNA-seq and Perturb-seq and 17- to 41-fold for Flex, with high concordance with Cell Ranger. As the MorPhiC consortium’s production processor, the suite has been tested and updated against large, modern production datasets. All of this was accomplished with AI-assisted development, which not only built the 183,606 new lines of C/C++ but has kept them maintainable through eight months and 25 releases in production. The result is an open transcriptomics processor, used in production at consortium scale, that humans and agents can run with a single command, and that a community using AI can now maintain and extend.

## 5 Methods

### 5.1 Benchmarking datasets

All benchmarking datasets are listed in Table 1. Two MorPhiC data production centers contributed data: the Memorial Sloan Kettering Cancer Center (MSK) and the Jackson Laboratory (JAX). STAR-core scRNA-seq parity was validated on the public 10x PBMC 10K (3’ v3) dataset (638.9 million read pairs) against a Cell Ranger 9.0.1 run made for this work, because the outputs published with that dataset come from Cell Ranger 3. Upstream STAR 2.7.11b under the community-optimized STARsolo parameters (CellGENI ^29^) was run on the same data as the baseline. The same three-way comparison was repeated on two further public datasets: a glioma cell line (GEO GSE261042, SRA run SRR28247645; 3’ v3, 117.0 million read pairs) and an organoid (GEO GSE316010, SRA run SRR36749062; GEM-X 3’ v4, 301.6 million read pairs), each against a Cell Ranger 9.0.1 count run made for this work (Supplementary Table S1). STAR-Perturb was benchmarked on two datasets. The A375 CRISPR dataset has 11 features, and Cell Ranger assigns a guide to 1,083 of its cells. The MSK 30-KO Perturb-seq dataset on ES cells, produced by the Huangfu Lab at MSK, combines gene expression, 30 CRISPR guides and 245,979 LARRY lineage barcodes over 33,226 cells. STAR-Flex was benchmarked on three datasets, all processed with GRCh38-2024-A annotations and probe set v1.1.0. The JAX SC2300771 Flex dataset (4 biological tags, 8 lanes, 2.011 billion read pairs) was compared against Cell Ranger 9.0.1. The public GSE325982 pool (four single-tag samples, 1.121 billion read pairs) was compared against our own Cell Ranger 9.0.1 run, made with the same settings as the other two datasets. The public 10x 320k scFFPE GEM-X dataset (sixteen tags pooled in pairs into eight samples, 7.303 billion read pairs) was compared against our own Cell Ranger 9.0.1 run. STAR-SLAM was benchmarked against GRAND-SLAM on the publicly available GRAND-SLAM 100K-read human fixture, which carries reference new-to-total RNA ratio (NTR) values. No SLAM-seq speed comparison was made, for the reasons given in Results.

### 5.2 Benchmarking metrics

For deterministic components—adapter trimming and coordinate sorting—parity was measured as exact identity: trimmed FASTQs were hashed against fixed Cutadapt output to confirm byte-identical files, and BAM records were compared after blocked GZIP (BGZF) decoding and coordinate sorting. Gene-level parity was assessed on raw (untransformed) per-gene count totals over the cells called by both tools, over every gene present in both annotations, by Spearman correlation (tied ranks averaged) and Pearson correlation. We report the Spearman correlation alongside the Pearson because a Pearson correlation, or any correlation restricted to well-expressed genes, is dominated by the most abundant genes and hides differences in low-count genes, which is where counting rules differ most. Bulk quantifications were compared the same way on the read counts assigned to each gene and each transcript (the `NumReads` column of Salmon’s output, which TranscriptVB writes in the same format), on the no-Y-removal benchmark. Cell-level parity was assessed by two measures on the cells called by both tools: cell Pearson, the Pearson correlation across cells of each cell’s total UMI count, and mean per-cell Pearson, the Pearson correlation across genes of one cell’s log(1 + *x*)-transformed counts in the two outputs, averaged over cells. The latter is taken over genes with *≥*20 counts in both outputs and detected in *≥*1% of shared cells, and cells whose counts do not vary are skipped. For the multiplexed Flex datasets a cell is identified by its 16-base barcode and its sample tag, and every measure is computed pooled and per sample as described in Results (Supplementary Table S3). Where STAR-Flex and cyto are compared with each other rather than each with Cell Ranger, the measures are computed on the cells all three tools call, so that both are scored on the same cells (Supplementary Table S8). The concordance script is provided in the benchmark repository. Cell-calling overlap was assessed by the Jaccard index ^47^ of the two called-cell sets *A* and *B*, *|A ∩ B|/|A ∪ B|*. Feature barcode parity was assessed by per-guide Pearson correlation on unique molecular identifier (UMI) counts, per-cell total UMI Pearson, exact called guide-set match rate, and directional containment metrics (the fraction of Cell Ranger calls recovered by STAR, and vice versa). SLAM-seq parity was assessed by Pearson and Spearman correlation on NTR and raw conversion fractions (k/nT) at multiple read-count thresholds (*≥*20, *≥*50, *≥*100). Runtime comparisons used the wall-clock time of the whole process from `/usr/bin/time -v` for every tool, with sequential runs to avoid contention and the page cache dropped before each run. Cell Ranger 9.0.1 was run with BAM output disabled and secondary analysis disabled (`create-bam,false` and `no-secondary,true`), with the same thread count as STAR Suite, and timed to completion, because its output directory is delivered only when the pipeline finishes. Where the same work needs two Cell Ranger runs—the MSK 30-KO dataset, one run per feature library—the two wall times are summed. cyto was release 0.4.7 in every run, called by absolute path with its version recorded in the run log, so that no run in the panel mixes releases. The time at which Cell Ranger writes the per-sample matrices, shortly before it completes, is given in Supplementary Table S5 for reference. All timings in Table 2 except those of the 320k dataset were measured on one machine (Intel i9-13900KF, 24 cores running 32 threads, 128 GiB of memory) with inputs, outputs and scratch on the same local NVMe device, for STAR Suite, cyto and Cell Ranger alike. The 320k dataset was measured for all three tools on an AWS instance limited to that machine’s size—an m6id.16xlarge (Intel Xeon Platinum 8375C) at 32 virtual CPUs and 128 GiB of memory, with two local NVMe devices in RAID0, Ubuntu 24.04—because its Cell Ranger run needs about 3 TB of scratch and most of a day; there Cell Ranger ran with 32 cores and 120 GB of memory and cyto under a 120 GiB memory limit. The spinning-disk comparison and the instance matched to it are described in Supplementary Note 2. The high-water mark of disk use was recorded by sampling every 5 s during each run.

Parity metrics are computed on the full, non-Y-removed read sets, so STAR Suite and the reference tool are compared on identical data. Y-chromosome removal, which the consortium’s KOLF2.2J line requires (Results), enters only the bulk speedup benchmark, where it is timed both integrated in STAR Suite and performed by the original external tools. The single-cell benchmarks leave it out on both sides, because Cell Ranger does not remove it and a step only STAR Suite performs would inflate the speedup without reflecting what the community runs. For the same reason the STAR Suite single-cell runs do only the work the Cell Ranger runs do: they count genes only (--soloFeatures GeneFull), without the Velocyto spliced and unspliced matrices that Cell Ranger does not produce, and write no BAM (--outSAMtype None), as Cell Ranger was run without BAM output.

### 5.3 Flex benchmark conditions

Cell Ranger’s probe set can be used in full or restricted to the probes it marks as included. Every run of every tool reported here used the default, the included probes only, on all three datasets, and cyto’s probe-to-gene table was built from the same probe set CSV. GSE325982’s submitters processed their deposit with filtering off; we ran Cell Ranger that way as well, reproducing their matrices byte for byte, and that run is compared with the default one in Supplementary Note 4 (Supplementary Table S11).

The 320k dataset has two tags per sample, whereas the JAX and GSE325982 datasets have one. Cell Ranger and STAR-Flex call cells per biological sample (Cell calling), whereas cyto filters cells for each tag separately and the per-tag matrices are then concatenated by sample, as cyto documents ^48^ and its maintainer recommends ^49^.

Three input formats were benchmarked for Flex. The JAX and 320k datasets are delivered as BGZF and were read through the in-process parallel reader (STAR-core). The European Nucleotide Archive re-compresses what it distributes as a single gzip stream, so GSE325982 was read instead through STAR’s `--readFilesCommand` route with rapidgzip 0.16.0 (`rapidgzip -d -c -P 8`), an external program that is not part of STAR Suite. Because cyto is built around CBQ (BINSEQ) input, both tools were also benchmarked on that format; the CBQ files were produced once with the BINSEQ encoder, and the conversion time is excluded from every timing, for both tools.

For every timed run, each input, output, temporary and spill location was resolved to its physical device before the run began, and a run was rejected if any of them fell on a device other than the one intended. Without this check a tool that writes little can appear to run on slow storage while its heaviest writes go elsewhere, since STAR-Flex’s bucket spill directory, temporary directory and output prefix are separate settings. Disk use is reported for every tool as the peak of that sampling, the high-water mark above the pre-run baseline, because that is the capacity a site must provide. It is not the volume written over a run, which is larger for a tool that deletes intermediates as it goes: Cell Ranger runs its stages as separate processes and removes their outputs as the run proceeds, so on the JAX dataset its pipeline accounted for 2.10 TiB written against a 564 GiB peak. Supplementary Table S9 reports the peaks.

### 5.4 Software architecture

STAR Suite extends STAR 2.7.11b by organizing the codebase into module-focused directories (Fig. 1) while keeping a single shared copy of the STAR core in `core/legacy/`. New functionality is organized into three directories: `core/features/`, which holds shared overlays (VB/EM quantification, Y-chromosome removal, BAM sorting, feature barcode tools, and `libscrna`); `flex/`, the Flex pipeline; and `slam/`, the SLAM-seq module. Table 7 summarizes these components and their roles. A top-level Makefile builds the STAR binary with every module compiled in (`make core`) and the standalone tools of each module (`make flex-tools`, `make slam-tools` and their counterparts; `make all` builds everything). The result of the core build is a single STAR binary; when new flags are not invoked, behavior is identical to upstream STAR 2.7.11b. The C/C++ line counts reported for STAR Suite—a 28,228-line upstream STAR 2.7.11b base, 183,606 lines added, and 211,834 total at release v1.9.5—are produced by `tools/count cpp lines.sh`, which counts the four source trees the binary is built from (`core/legacy/source`, `core/features`, `flex/source` and `slam/source`) against a specified upstream STAR tag. It skips symlinks and excludes the vendored htslib, opal and klib libraries and the PCG random-number headers, so reviewers can rerun it to reproduce these figures. The subsections that follow describe the configuration each benchmark used rather than every mode the tools provide: the full option set is documented in the repository, the exact flags of every run are recorded in STAR-suite-provenance, and the scripts that issued them are in STAR-suite-paper.

**Table 7.**
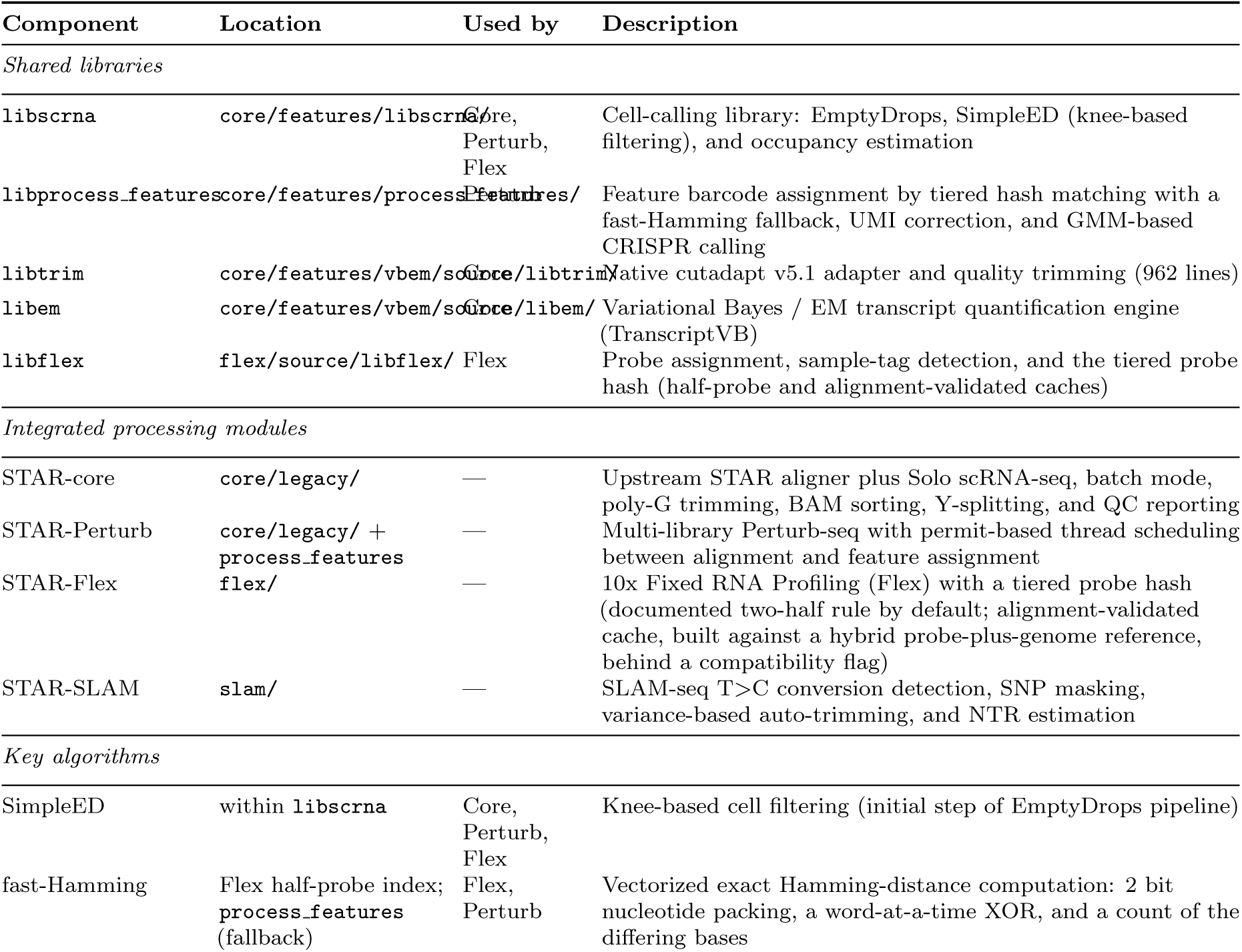
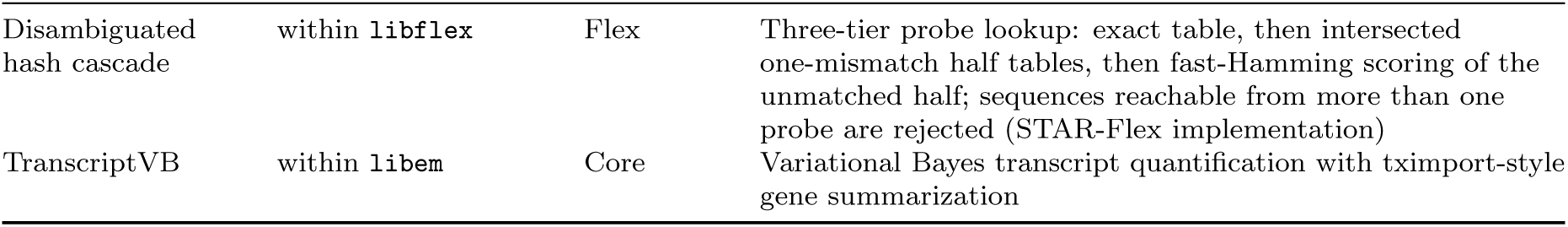
STAR Suite component glossary: internal modules and their roles. Component is the library or module name, Location its directory in the repository, and Used by the modules that call it; an em dash marks a module that is itself one of the four and so is not called by another. Description summarizes what the component does.

STAR Suite adds one dependency to upstream STAR’s, OpenSSL’s libcrypto, a standard system library, so it keeps STAR’s deployment model of a self-contained executable, including static binaries, which helped make STAR the standard transcriptomics engine. All new functionality is implemented in C/C++ using the standard library, the third-party headers already present in upstream STAR (htslib for BAM I/O^50^, opal for SIMD alignment^51^), and two small vendored header-only libraries (PCG random number generation^52^, 3,623 lines, and the klib khash, 617 lines); none of the vendored code is included in the line counts above. libcrypto supplies the SHA-256 checksums in the Flex reference manifest and the TranscriptVB shard sidecars. If STAR compiles and the OpenSSL headers are present, STAR Suite compiles. Binary-only deployment matters most on HPC clusters, where administrators install software on behalf of their users and every new dependency is a request that has to be justified and scheduled.

### 5.5 STAR-core: components shared across modules

#### Input: parallel BGZF FASTQ and binary CBQ

In BGZF, the blocked gzip in which sequencers deliver FASTQ (Results), each compressed block is independently decodable and carries its size in a `BC` extra subfield. Decompressing BGZF blocks in parallel is well established (htslib’s multithreaded reader does it for BAM). STAR Suite detects BGZF from the first file header `(--readFilesBgzfMode`, `auto` by default, with `off` and an explicit range mode as alternatives) and reads each file through a pool of inflate workers that process each batch on demand by hopping the declared block sizes (Fig. 2). Plain (non-BGZF) gzip continues through the standard stream unchanged, and this reader does not use or modify the vendored htslib.

The experimental binary-sequence input mode `(--readFilesType Binseq)` reads Arc Institute BINSEQ^21^ files (.cbq) natively; its current scope and unsupported cases are documented in docs/EXPERIMENTAL BINSEQ INPUT.md. The format is also compact for read sets that a consortium retains and periodically reprocesses: the 320k dataset is 434 GB as original FASTQ and 278 GB as CBQ.

#### Adapter trimming

STAR Suite implements the Cutadapt ^53^ trimming algorithm natively in C++ `(--trimCutadapt Yes`), removing the decompression and recompression of FASTQ files. We first validated it to byte-level parity with Trim Galore ^54^ (cutadapt 3.2), confirming the implementation was correct, then updated the default to the current Cutadapt v5.1. A compatibility mode `(--trimCutadaptCompat Cutadapt3`) retains the older behavior for byte-level agreement with legacy Trim Galore pipelines. The trimming module (libtrim/, 962 lines) is a general-purpose feature usable with any STAR workflow.

#### BAM sorting

By default, STAR sorts by coordinate with a bin-based scheme: during mapping it writes every alignment to one of a fixed set of genomic-coordinate bins as per-thread temporary files, then loads each bin in full into memory to sort it and concatenates the results. Spreading reads across bins helps only to the extent the bins are occupied—the memory requirement is set by the largest bin, and the temporary files hold the entire alignment set on disk however the reads distribute. The alignment-based Flex reference breaks that scheme: it adds a pseudo-chromosome for every probe, more than 50,000 sequences of 50 bases each, and with that many reference sequences the bin-based sort exhausted both memory and disk on production-scale runs and never completed. STAR Suite adds an alternative backend `(--outBAMsortMethod samtools)` modeled on `samtools sort`^55^: it accumulates records in a bounded in-memory buffer `(--limitBAMsortRAM)`, spills a sorted run to disk only when the cap is exceeded, and produces the final ordering by a k-way merge of those runs (the algorithm is reimplemented natively; no samtools code is used). Because memory is capped irrespective of how coordinates are distributed and records reach disk only under memory pressure rather than unconditionally, the alignment-based Flex routes, which are retained behind `--flexLegacy` yes, use this backend whenever coordinate-sorted output is requested.

#### Y-chromosome splitting

Reads with and without a chromosome-Y alignment are written to separate BAM or FASTQ files (--emitNoYBAM, --emitYNoYFastq) while the alignments are produced, so that Y-stripped FASTQ can be redistributed without a further pass over the BAM.

#### Cell calling (libscrna)

STARsolo already uses its own EmptyDrops ^30^-based cell calling, but it assumes a single sample per run, whereas Flex multiplexes several samples and must call cells separately for each. Rather than maintaining two callers, we consolidated the logic into one shared C library (`libscrna/)`—providing EmptyDrops, SimpleED, and occupancy-based filtering—used by both the non-Flex scRNA-seq and Flex pipelines; the two branches share the code and differ only in the settings below. The non-Flex branch estimates `recovered_cells` adaptively by bootstrap (100 samples), our own reimplementation of the automatic estimation that Cell Ranger 9 documents in place of a user-supplied `expect-cells`^56^, with 10K Monte Carlo simulations for EmptyDrops and Benjamini–Hochberg false discovery rate (FDR) correction ^57^. The Flex branch calls cells per biological sample: the tags belonging to one sample are grouped into one model while each cell keeps its full barcode-plus-tag identity, so the pooled pairs of tags in the 320k dataset are called as one sample each, as Cell Ranger does. It uses a fixed ambient rank window (Cell Ranger’s 45,000–90,000 barcode ranks, scaled by the sample’s tag count), 100,000 Monte Carlo simulations and Benjamini–Hochberg FDR 0.01, and an occupancy step after all samples are called removes barcodes seen in more sample tags than a fitted zero-truncated Poisson allows. Independent samples are called concurrently under the same shared thread budget as the rest of the run. Because the Monte Carlo draws are partitioned across bootstrap streams, the stream count is part of the reproducibility contract (Supplementary Table S6, class 1). On small or downsampled samples the library skips the Monte Carlo tail rescue and keeps only the high-count calls, which tracks Cell Ranger more closely than rescuing cells from a sparse ambient tail.

#### Permit-based thread scheduler

Alignment and feature assignment run in one process, and their threads are governed by a single pool of permits `(--dynamicThreadInterface 1`), by default one permit per configured thread. A thread does not migrate between workloads; instead each unit of work—one chunk of reads to align, one batch of reads to match against the feature library—must hold a permit while it runs, so the number of threads each workload has at any moment follows demand. When alignment stalls on input, its threads release permits that feature assignment acquires, and the reverse. Each workload can be given a borrowable floor, a number of permits it is guaranteed whenever it has waiting work, with the surplus shared freely; waiters are admitted in arrival order so that no workload can starve; and the pool size can be retuned during the run from measured processor idleness. Every release of a permit returns telemetry (wait time, units of work, bytes) for its workload, written to the log on request. The Flex cell caller draws on the same budget when it calls independent samples concurrently. In STAR-Flex the BGZF inflate workers also hold permits from this pool: a rate controller proposes one-permit changes to the decoder’s share from queue pressure and keeps a change only if the measured throughput of the consuming threads improves, so decompression takes threads only while the processors can use its output. The same contract governs the ATAC workload of the companion Chromap Suite, so a joint RNA and ATAC run shares one pool across three workloads.

### 5.6 Variational Bayes transcript quantification

STAR Suite performs Salmon-style ^28^ transcript-level abundance estimation—Variational Bayes by default, with an EM alternative—natively within the aligner `(--quantMode TranscriptVB; libem/, 26 files)`, avoiding a separate Salmon run. It also provides library-type auto-detection, fragment-length distribution modeling, and tximport-style^58^ gene-level summarization `(--quantVBgenesMode Tximport)`, and generating the transcriptome FASTA during index building `(--genomeGenerateTranscriptome Yes)` removes the separate gffread or rsem-prepare-reference step. The same EM code backs the SLAM-seq module’s conversion-rate mixture model.

### 5.7 Multi-sample execution and reference construction

`--batchMode 1` processes several samples in one invocation with the genome index loaded once. The samples’ FASTQ files are given as comma-separated lists in `--readFilesIn`, one list per mate, and each sample’s outputs are routed to its own subdirectories (`alignments/, qc/, counts/, y separated/` `<sample>/)` under the output prefix. A sample that fails stops the run, or on request `(--batchOnError respawn)` STAR relaunches itself on the remaining files, up to `--batchMaxRetries` times. Batch mode is available for bulk and SLAM-seq runs. STARsolo, two-pass mapping and junction-based read filtering are not available in it.

`--autoIndex` Yes, with `--runMode genomeGenerate`, builds the index in the genome directory unless one already exists. When no FASTA, GTF or URL is supplied it downloads the Ensembl GRCh38 primary assembly and the GENCODE annotation for the Cell Ranger reference release named by `--cellrangerRefRelease`, either 2024-A with Ensembl 110 and GENCODE 44 or 2020-A with Ensembl 98 and GENCODE 32, into a cache under the genome directory, and verifies their checksums. It can also format them as Cell Ranger does `(--cellrangerStyleIndex` Yes: chromosome renaming, biotype filtering and version stripping) before building the index`. -genomeGenerateTranscriptome` Yes adds the transcriptome FASTA (Variational Bayes transcript quantification).

### 5.8 scRNA-seq processing

The single-cell changes are to STARsolo’s read clipping and counting, not to alignment. Each is a flag that our Cell Ranger comparisons set.

#### Poly-G clipping

Two-color sequencers (NovaSeq, NextSeq) call the absence of signal as G, so a read that runs off its insert ends in a poly-G tail. Upstream STAR’s Cell Ranger-style clipping removes only poly-A. A poly-G read aligns perfectly, uniquely and at full mapping quality to a genomic G homopolymer. In GRCh38 a run of about 90 bases lies inside the lncRNA LINC00486, the gene that is over-counted when such tails are left in place (Results). `--clip3pPolyG yes` clips a 3*^′^* run of G with the scan STAR uses for poly-A, and the default `auto` turns it on whenever `--clipAdapterType CellRanger4` is set. Working in from the 3*^′^* end, a G scores +1 and any other base *−*2. The longest prefix whose running score is at least 70% of its length is taken as the candidate clip, the scan stops early once the score has fallen more than 27 below the length examined, and the candidate is clipped only if its score reached 20. Because a mismatch costs twice what a match earns, the 70% condition admits at most one non-G base in ten, so an accepted tail is at least 90% G and at least 20 bases long. Reads shorter than 20 bases are not clipped, and the longer of the poly-A and poly-G clips is taken.

#### Multimapper rescue

STARsolo counts a read that maps equally well to several places only under the redistribution rules of `--soloMultiMappers,` which was set to Unique (such reads are not counted) in every benchmark here. `--soloCrMultimapRescue` yes adds a rule, inferred from matched Cell Ranger outputs (Section 2.8), that resolves such reads before counting. Each equal-best alignment is classified as exonic, intronic or unannotated with respect to the countable genes, where pseudogene biotypes are not countable except for the immunoglobulin and T-cell receptor pseudogenes. A read with exactly one exonic candidate is assigned to it. Failing that, when the intronic fallback is on, a read with no exonic and exactly one intronic candidate is assigned to that gene; the fallback `(--soloCrMultimapRescueIntronic)` is on by default whenever the matrix written in Cell Ranger’s layout is GeneFull. Any other pattern is left unresolved. Excluding pseudogene copies from the countable set is what brings highly expressed genes with many pseudogene copies to Cell Ranger’s counts (Results). With GeneFull a second rule handles a small gene lying inside another gene’s intron: an alignment that overlaps several gene bodies, only some of which it matches exonically, is counted for the exonic genes only.

#### Intron-inclusive counting and the Cell Ranger-layout matrix

Cell Ranger 7 and later count intronic reads. The suite matches this by requesting `--soloFeatures GeneFull` and naming Gene-Full as the matrix written in Cell Ranger’s layout `(--soloCrGexFeature genefull`; `auto` writes Gene when it is present and GeneFull otherwise). Barcode correction, UMI filtering, UMI deduplication and cell calling use STARsolo’s Cell Ranger-style settings (`1MM_multi_Nbase_pseudocounts, MultiGeneUMI_CR, 1MM_CR, EmptyDrops_CR; Cell calling).` The preset `--defaultCrCompat yes` applies these defaults for the Perturb-seq path together with its feature-calling defaults (Feature calling); it does not enable poly-G clipping, which the benchmark commands set explicitly.

#### On-chip multiplexing

GEM-X OCM libraries carry the sample in bases 8 and 9 of the 16-base cell barcode. With `--ocmMultiEnable` in composite-barcode mode `(--ocmMultiBarcodeMode flex)`, STAR appends an eight-base sample tag to each cell barcode before barcode correction, so that correction and cell calling never cross sample populations, and writes per-sample raw and filtered matrices in Cell Ranger’s multi layout, calling cells per sample after the split.

#### Velocity matrices

When `--soloFeatures Velocyto` is requested, STAR writes raw and filtered Velocyto-compatible matrices under `outs/`. The spliced, unspliced and ambiguous layers are built from the same clipped reads and the same resolved alignments as the gene matrix, because clipping and multimapper rescue run before the layer classification, and they take their corrected barcode, corrected UMI and keep-or-drop decision from the gene feature. The layers keep STAR’s Velocyto semantics and are not constrained to sum to the GeneFull matrix.

### 5.9 Feature barcode assignment

Feature barcode assignment matches each read to a feature in the reference library—a CRISPR guide or a lineage barcode—producing the per-cell feature UMI counts; the motivating scale problem and the speed and concordance benchmarks are in Results (STAR-Perturb). The feature barcode engine (`process_features, in core/features/process features/)` is used in integrated STAR-Perturb and in standalone feature-calling tools. It accepts a standard Cell Ranger feature reference CSV as input. For multi-library experiments, chemistry can be set explicitly or auto-detected per library to support mixed TRU/NXT inputs.

Matching is tiered. Most reads match a feature exactly, or within a small Hamming distance, at or near the expected offset, so the engine first looks the read up in hash tables of the features and of every variant within one and within two mismatches (Fig. 3b). The tables are built once per library; a variant that more than one feature could have produced is marked ambiguous at that point, and a read that lands on it is left unassigned. Enumerating variants becomes intractable as the distance grows, since their number grows combinatorially, and the tables are skipped when their estimated size would exceed the memory budget.

Behind the tables sits fast-Hamming, the vectorized exact Hamming-distance search that STAR-Flex also uses to decide probe matches: sequences are packed two bits per nucleotide. The bitwise XOR is used to compare up to 64 bits at a time. Words that are equal are skipped without counting, and the comparison is abandoned as soon as the running total passes the mismatch ceiling, so an unrelated barcode is usually rejected on its first word. At each 2 bit position the XOR gives a result of zero only if the two bases are the same; otherwise it gives either one or two set bits. Our original `process_features` implementation used a lookup table to convert these bit patterns into a Hamming distance 8 bits at a time. This is most efficient when long sequences are being compared and is completely independent of the underlying hardware and is the approach we use for feature detection where longer sequences might be compared. As an alternative we can use bit operations to fold the two bit results into the lower bit. There are four possible 2 bit outcomes after the XOR: 00, 01, 10 and 11. If we shift right by one, the four outcomes become X0, X0, X1 and X1, where X is the bit shifted in from the neighbouring position. We then OR against the original string. X0 — 00 = X0, X0 — 01 = X1, X1 — 10 = 11, X1 — 11 = 11. After this transform the lower bit in each case is 1 whenever there is a difference and 0 when the two bits were identical, that is, when the bases match. We then filter out the higher bit by masking with 01 at each position. This gives us 00 when the original bits were identical and 01 for everything else. The total number of bits in the result is the Hamming distance. We count those bits with the compiler’s own intrinsic, `__builtin_popcountll`, which GCC compiles to the hardware population-count instruction only when the target is told that the instruction exists (`-mpopcnt`, or a baseline of x86-64-v2 or above), and otherwise to a portable software routine. The instruction has been present on most x86 processors since about 2008, but a precompiled binary cannot assume it: built that way it would fault on any machine without it. Our releases are therefore built for the generic baseline and take the software routine, and it is a user building from source who can turn the instruction on. For longer sequences, the compatible method is slower than the lookup table but the hardware method is faster.

For Flex we knew we were dealing with exactly 25 bases and the LUT and popcount method gave almost identical results, so we went with the popcount method which would be faster if the hardware was available and enabled.

For feature barcodes it is the fallback, engaged at mismatch ceilings above two or when a library’s variant tables are not built; it is general enough to scan even the full 245,979-barcode LARRY library on the MSK dataset without approximation or indexing heuristics. The combination keeps the speed of a hash search with the guarantee of finding the best Hamming distance whenever the search is needed. For the MSK Perturb-seq we used a one-mismatch ceiling. So while fast-Hamming was developed for feature processing, in this paper, it is in STAR-Flex where fast-Hamming is actually used. Feature UMIs are then deduplicated per cell and feature by merging UMIs within one mismatch of each other into connected components, whereas the gene-expression library of the same run is collapsed by STARsolo’s Cell Ranger-style rule (scRNA-seq processing).

### 5.10 Feature calling

Whereas assignment matches reads to features and tallies their UMI counts, feature calling is the per-cell decision of which of those features a cell genuinely carries—the CRISPR guide it received, or the lineage barcode marking its clone—separating true feature signal from background. When processing `--pfMultiConfig` with CRISPR Guide Capture features, STAR runs it automatically after EmptyDrops filtering. For CRISPR guides the caller is a Gaussian mixture model that sets per-feature UMI thresholds for assigning calls to cells, as Cell Ranger documents `(--crGuideCaller);` its minimum UMI threshold `(--crMinUmi)` is assay dependent and was set to 10 for the A375 benchmark and 2 for the MSK guides. For lineage barcodes, where the question is which one barcode marks a cell’s clone, the caller is the production dominance rule: on the filtered lineage matrix, the barcode with the highest UMI count is called when its count is strictly greater than the second-highest, and a tie leaves the cell unassigned. It runs inside the STAR pass (`star_feature_caller=dominant` in the multi-library configuration). On the MSK benchmark it gave the same call as the production script for every one of the 26,521 cells shared with the released production matrix, 26,017 identical assignments and 504 cells unassigned by both (record `runs/msk_30ko_es_v195` in `morphic-provenance`). This calling validation was separate from the timed P02 configuration, which left the opt-in dominant caller disabled. Output files `(crispr_analysis/)` match the Cell Ranger-compatible layout (Standalone utilities).

### 5.11 STAR-Flex implementation

Probe assignment uses one of two cache types, which differ in how they settle an ambiguous match. The first is the alignment-validated hash cache whose rationale and history are given in Results (STAR-Flex): built against a hybrid reference of the genome plus one pseudo-chromosome per probe, it retains a single-mismatch variant only when alignment maps it to its probe rather than to the genome, so alignment acts as an oracle against genomic background.

The second, the half-probe cache, removes the check against the rest of the genome. Each 50-base probe is divided into two 25-base halves. The cache stores a table of the exact probe sequences and, for each half, a table of every sequence within one mismatch of that half, listing the probes each can come from. Reads pass through three tiers. A read that matches a probe exactly is assigned directly. Otherwise both of its halves are looked up and their probe lists intersected, and a read whose two lists have exactly one probe in common is assigned to it, while a read whose lists have more than one probe in common is rejected as ambiguous. When only one half is found and it identifies a single probe, the read is anchored to that probe and its other half is compared with the same probe by fast-Hamming (Feature barcode assignment); the read is accepted when the anchoring half’s mismatches (at most one) and the other half’s together number at most ten. A read is rejected when neither half is found, when its two halves point to different probes, or when the anchored comparison exceeds ten mismatches. At most one mismatch is permitted per half, which keeps the tables small enough to query for every read.

The half-caches are disambiguated when they are queried, not while they are built. We vary the sequence at each position to generate all possible single-mismatch variants of each half and check them against a hash table of variants. A variant that has not been seen before is added, recorded as a variant of the probe that generated it; a variant that another probe has already generated keeps both probes. This is where the half tables differ from the feature-barcode tables of the previous subsection, which enumerate the variants of a whole feature and mark as ambiguous, when the table is built, any variant that more than one feature could produce. A half table cannot settle a probe on its own, and it answers two different questions: intersected with the other half’s list it identifies the probe, and taken alone it supplies the anchor probe whose other half is then scored by fast-Hamming. A variant that two probes share is therefore kept rather than discarded, because the other half usually settles it; for the JAX probe set the two tables hold 8,296,122 half keys, of which 38 are shared by more than one probe. The cache depends only on the probe set and not on the reads, so it can be pre-generated (2 min 18 s on 32 threads for the JAX probe set; Supplementary Note 1) and loaded like a reference; its construction is therefore not part of the run times reported here.

Cell-barcode and UMI resolution is independent of sample-tag resolution: the two are determined separately, so a read whose tag cannot be resolved still yields a cell barcode and UMI. That independence is what allows tag detection to run inline within the single read-processing stream rather than as a separate pass over the data.

Sample tags are resolved against the probe-barcode table supplied with the assay, by exact lookup first and then, unless strict matching is requested, by a single-mismatch tier over that same table. The tolerance is a parameter `(--soloSampleTagMismatch, default 1)` restricted to zero or one, and `--soloSampleStrictMatch` reduces it to exact lookup alone; an eight-base window containing an uncalled base is rejected before either lookup. Each run reports its exact, single-mismatch and ambiguous tag counts separately. Probe windows take the opposite route: a single uncalled base among the 50 is resolved by substituting each of the four bases in turn and putting all four through the tiers above `(--soloProbeMismatch, default 1`). The read is assigned only if the substitutions that resolve agree on one probe and none of them is a denial. 10x documents that the barcode table is used, but the single-mismatch tolerance was one of the behaviours we discovered by examining the outputs.

Every Flex benchmark reported (JAX, GSE325982 and 320k) was processed with the half-probe cache. The alignment-validated cache, and full alignment of cache misses, remain available behind a compatibility flag `(--flexLegacy yes)`. Supplementary Note 3 compares the two probe-assignment routes.

The full Flex pipeline runs as a single in-process stream: probe assignment against the half-probe cache (Fig. 3a), inline sample-tag detection, single-mismatch pseudocount-based cell-barcode correction, connected-component UMI correction, and cell filtering grouped by biological sample through EmptyDrops, SimpleED, and occupancy-based refinement (see Cell calling). UMI correction uses the same one-mismatch connected components as feature barcode assignment. A cell is the composite of the corrected 16-base cell barcode and the 8-base sample tag; UMI correction, deduplication and counting are performed within that composite and one gene, so UMIs are never merged across samples and reads of one GEM barcode carrying different tags contribute to different cells. No per-barcode sample call is made.

### 5.12 STAR-SLAM implementation

STAR-SLAM performs T*>*C conversion detection, SNP masking, auto-trimming, and gene-level NTR estimation within the aligner’s read-processing loop; the production problems that motivated moving this analysis into the aligner—logic drift, paired-end counting for DESeq2, and per-sample variant masking—are set out in Results (STAR-SLAM). For each aligned read, genomic positions are inspected for T*>*C conversions (sense strand), with positions matching known SNPs excluded. The NTR is estimated per gene using a binomial model with EM inference, computing Maximum A Posteriori (MAP) and mean estimates with 0.05/0.95 quantile credible intervals from Beta posterior parameters. Gene-level results are written by default in GRAND-SLAM’s table format `(--slamGrandSlamOut, <prefix>SlamQuant.grandslam.tsv),` carrying its column names for read count, the posterior mean, MAP and 0.05/0.95 quantiles, the Beta parameters, conversions, coverage and gene length, so that tools written against GRAND-SLAM’s output read STAR-SLAM’s unchanged.

SNP masking normally uses an external mask of known variant positions, a bgzip-compressed, tabix-indexed BED derived from the sample’s VCF `(--slamSnpMaskIn)`—the route we use in practice, since variant calls are available for most samples today. To mirror GRAND-SLAM, which instead estimates SNP sites from the SLAM-seq data itself when no variant calls are supplied, STAR-SLAM also provides an internal auto-detection mode `(--slamSnpDetect 1):` a Kneedle-style knee detection ^59^ on the per-position mismatch-fraction distribution that builds a histogram of `mismatches/coverage` at positions with *≥*10 coverage, applies `log1p` normalization, and takes the bin maximizing distance from the diagonal. Guardrails require *≥*1,000 eligible sites and knee strength *>*0.02 and clamp the result to [0.10, 0.60], with fallback to GRAND-SLAM/GEDI’s default of 0.22. This path exists for GRAND-SLAM parity rather than routine use.

At read ends, the baseline T*>*C transition noise is far more variable—chemical modification artifacts distort the conversion rate there, and because these artifacts do not track Phred quality scores, quality-based trimming leaves them in. Variance-based auto-trimming `(--autoTrim variance)` locates the reliable region from this variability: it computes the standard deviation of T*>*C conversion rates across read positions, fits a piecewise linear regression (2, 3, or 4 segments, selected by the Bayesian information criterion) to the smoothed standard-deviation curve, and takes the breakpoints as the reliable-region boundaries. Trim values can be computed once from the first input file and applied globally `(--trimScope first, default)` or independently per file `(--trimScope per-file).`

The reference GRAND-SLAM implementation, in the GEDI platform, counts reads and resolves multi-gene overlaps using its own conventions, which differ from STAR’s native counting. To reproduce its outputs—for parity testing and for users migrating existing GRAND-SLAM workflows— a GEDI compatibility mode `(--slamCompatMode gedi)` switches four behaviors to match: (i) intronic classification of unspliced reads overlapping introns of multiple transcripts; (ii) lenient overlap acceptance (*≥*50% exon overlap + splice junction concordance); (iii) read-level overlap-gene weighting (weight = 1/nTr/geneCount); (iv) configurable position filtering for paired-end (PE) overlap and trim guards. Each compatibility feature can be overridden on its own, so one behavior can be changed without affecting the others.

### 5.13 Standalone utilities

In addition to the main STAR binary, STAR Suite provides standalone tools built from the same libraries as the integrated pipeline, so that a module’s outputs can be recomputed without repeating the work that produced them. `Slam_requant` carries the SLAM quantification engine (SlamQuant, SlamSolver, SlamCompat, SlamVarianceAnalysis) and requantifies from the per-read dump a STAR-SLAM run writes, so that SNP masks, auto-trim settings and model parameters can be varied on a finished run without re-aligning it. `Run_flexfilter_mex` re-runs Flex cell calling (SimpleED, EmptyDrops, occupancy filtering) with different parameters on an existing Market Exchange (MEX) matrix. `assignBarcodes` exposes the lower-level feature-assignment engine, and `star_feature_call` takes FASTQ through feature barcode assignment, EmptyDrops cell calling, GMM-based feature calling and Cell Ranger-compatible output, with no alignment at any stage.

### 5.14 QC reporting

The trimming, SLAM-seq, and feature-barcode modules produce interactive QC reports in HTML and JSON format. Trim QC reports `(--trimQcReport)` include adapter match rates, trimmed length distributions, and quality score profiles. SLAM QC reports `(--slamQcReport)` visualize per-position T*>*C conversion rates, the auto-trim variance curve with fitted breakpoints, SNP detection histograms, and per-gene NTR distributions. Feature-barcode QC reports render feature-type (richness and co-expression) and feature-count heatmaps together with feature multiplicity and richness histograms. Producing them requires nothing beyond the binary itself (Table 4). The JSON holds the full contents of each report; the HTML pages draw their plotting library from a content delivery network, so rendering them in a browser needs network access.

### 5.15 AI-assisted development workflow

The “Human-Architect, AI-Implementer” cycle that produced the codebase, and the division of labour it rests on, are described in Results (Fig. 4; Table 6). The researcher worked with the agents through the IDE and MCP-assisted workflows, and through most of the development period the agents were not reliable enough to run the implement-and-test loop unsupervised, so their work was reviewed and approved at most stages. The design record is kept in the repository as it was written: architectural plans and per-feature design summaries under `docs/` and `plans/`, the runbooks and handoff notes that carried each task between the developer and successive AI sessions (155 of the 275 working documents at release 1.9.5), release notes for every release, and the benchmark records under `docs/benchmarks/`. It is the record an AI agent reads before continuing the work, which is why it is maintained: in this workflow, provenance is not documentation written afterwards but the medium through which maintenance happens.

### 5.16 Agent tooling and documentation

An MCP server^44^ gives AI agents structured tooling: discovery `(list_datasets, list_test_suites, find_docs, find_tests),` recipe resolution (the recipe-catalog tools, which also lock a resolved recipe), pre-flight validation `(preflight),` build management `(build_star, ensure_fresh_build),` script execution (run script with allowlisting, collect outputs), the workflow-parameter tooling described below, and hot-reload of the server configuration `(reload_config)`. All paths are validated against trusted roots. A job queue (1 concurrent, 10 queued) with timeout handling and process group cleanup prevents runaway executions.

The workflow-parameter tooling is a service that exposes a machine-readable recipe for each production workflow. A recipe is a typed definition of a workflow’s parameters (string, int, float, bool, enum, file, directory, string list), their CLI flag mappings, default values, cross-parameter constraints (mutual exclusion, dependency, positivity), rendering rules for deterministic flag ordering, and semantic stage metadata. STAR Suite serializes each recipe as YAML; the file format is a serialization choice rather than part of the definition. Agents interact with this service through five core tools: `list_workflows` and `describe_workflow` for discovery; `get_workflow_parameter_schema` to retrieve the full parameter contract; `validate_workflow_parameters` to check a parameter set against the schema including type coercion, constraint enforcement, and filesystem path validation; and `render_workflow_command` to produce a normalized argv array and shell-safe command string from validated parameters. This structured contract replaces ad hoc flag guessing with a validate- then-render pattern: agents discover the workflow, retrieve its schema, propose parameters, validate them, and render the final command—all without executing anything until the human approves.

To bootstrap schemas for new workflows, a `scaffold_workflow_schema` tool parses an existing shell script’s `case`/`while` flag-parsing patterns and `usage()` heredoc to extract parameters, infer types, and generate a draft YAML that can be refined and committed. A companion `validate_draft_workflow_schema` tool checks draft schemas against the Pydantic model before they are loaded. The schema format and authoring process are documented in `mcp_server/workflows/AUTHORING.md.`

The MCP server presents each recipe to AI agents as schema-validated tool calls and, within the same process, hosts a browser-based STAR Launchpad (`mcp_server/launchpad/)` that renders the same recipe as a web form for human operators (Fig. 6). The consortium’s own production recipes are maintained separately, in `morphic-recipes`, as declarative YAML for the deployments at MSK, UCSF and JAX.

Each run record in morphic-provenance holds the STAR Suite commit and binary checksum, the rendered commands, the complete processing scripts around STAR Suite with their commits and checksums, input and output inventories with checksums, transfer and delivery status, and pointers to dated release notes describing what each consortium member received. Records are append-only: a correction is added with its reason and the fields or files it changed. At the time of writing the repository holds 16 run records across ten projects and 12 dated deliveries. A message-only rewrite of the STAR-suite repository history on 13 September 2026 removed automatically inserted co-author trailers from 16 commits without changing any tree, author or date; the mapping from the earlier commit identifiers to the current ones is kept in the repository (https://github.com/morphic-bio/STAR-suite/blob/master/docs/COMMIT_ID_MAP_20260913.tsv) so that provenance records made before that date remain resolvable; source trees, release binaries and recorded binary checksums are unchanged.

STAR Suite’s own recipes and records form a general layer beneath the consortium’s. The suite’s reusable recipes—general workflows, open scripts, and scheduler, storage and site profiles—are maintained in `STAR-suite-recipes.` Its benchmark, parity, release and correction records are maintained in `STAR-suite-provenance`, in a fixed schema checked by a validator that also rejects local paths and host names; published records are never edited, and a correction is a new record naming the one it supersedes. The records behind this paper’s benchmarks are among them. Because those records carry no paths or command lines, the scripts that produced the paper’s benchmarks—the run drivers, the acceptance checkers, the campaign gates and the concordance analysis—are published separately in `STAR-suite-paper`, with the locations of our machines replaced by configuration variables so that the protocol is inspectable and the runs are repeatable elsewhere. Each STAR Suite release carries a checked copy of both repositories under `share/star-suite/`, with a manifest of their revisions and checksums that the release build verifies, so the recipes and evidence a release was validated against are available offline. The recipe resolver stacks catalogs—STAR Suite’s, then a site’s, then a project’s—so a consortium extends the general recipes rather than copying them, and project run records are kept apart from the release archive.

The repository is documented for AI agents as much as for people. `AGENTS.md` at the root is an orientation map—repository layout, build commands, key technical decisions (with links to detailed summaries), data locations, branching policy, and CI/CD configuration—and `tests/ARTIFACTS.md` catalogs the gold-standard datasets and benchmark runs with their locations. Together with the design record above, these let an AI agent, or a new human contributor, orient in the codebase and reproduce past work without hand-holding.

### 5.17 Availability and requirements

Project name: STAR Suite. Project home page: https://github.com/morphic-bio/STAR-suite. Operating system: Linux (x86-64 binaries and multi-architecture x86-64 and arm64 Docker images are published with each release). Programming language: C/C++ (the MCP server and Launchpad are Python). Other requirements: to run the released binary, zlib—which STAR already required— and OpenSSL’s libcrypto; to build from source, g++ with C++17 support, GNU make and zlib. The default `make core` links the Chromap Suite library for multiome runs, `make core-static` builds the statically linked STAR-only binary carried by the release tarballs, and `make core-portable` the dynamically linked STAR-only binary built by the Debian packages, whose packaging lists the build tools and development libraries it needs. License: MIT. Any restrictions to use by non-academics: none.

### 5.18 Testing infrastructure

The test suite is organized by module: Cell Ranger-compatibility parity tests, CRISPR feature calling integration tests, Flex smoke and multi-sample tests, SLAM fixture parity and E2E tests, Y-chromosome splitting tests, parameter regression tests, and TranscriptVB/Salmon parity tests. Continuous integration runs on GitHub Actions. Every pull request builds the suite in a container, verifies the pinned recipe and provenance snapshots and runs the Tier A smoke tests, which exercise the suite on small fixtures; pushes to the dev-release and master branches repeat those checks and publish container images (amd64 and arm64 for master). A version tag runs the release workflow. It builds the binaries in Ubuntu 22.04 and 24.04 containers, then packages them as tarballs against two glibc baselines, an installer bundle and Debian packages. Each artifact is validated on the systems it targets and the tarball and installer bundle are smoke-tested on 24.04; every artifact is then checksummed, the Debian source package is signed, and the multi-architecture images and the GitHub release are published with its notes. The fixture-heavy tests (Tier B) and the production-scale comparisons are run outside continuous integration before a tag is cut, and their results are recorded in the release notes and in STAR-suite-provenance.

## Declarations

### Ethics approval and consent to participate

Not applicable.

### Consent for publication

Not applicable.

### Availability of data and materials

STAR Suite (release v1.9.5) is available at https://github.com/morphic-bio/STAR-suite under the MIT License. Production-grade MorPhiC workflow recipes are maintained in a companion repository at https://github.com/morphic-bio/morphic-recipes, and per-run provenance records linking each consortium output to its exact STAR Suite invocation are maintained at https://github.com/morphic-bio/morphic-provenance. A snapshot of all three repositories is archived at https://doi.org/10.5281/zenodo.21632664, a concept identifier that resolves to the most recent archived version. STAR Suite’s general recipes and its benchmark and release records are at https://github.com/morphic-bio/STAR-suite-recipes and https://github.com/morphic-bio/STAR-suite-provenance, and each release includes a checked copy of both. The scripts that produced this paper’s benchmarks are at https://github.com/morphic-bio/STAR-suite-paper.

The GRAND-SLAM benchmark fixture dataset is publicly available at https://github.com/erhard-lab/gedi/wiki/GRAND-SLAM. The A375 10x CRISPR 5’ GEX dataset is publicly available from 10x Genomics (https://www.10xgenomics.com/datasets/1k-CRISPR-5p-gemx). The MSK 30-knockout Perturb-seq and JAX SC2300771 10x Flex benchmark datasets are MorPhiC consortium datasets under embargo pending consortium data release; they will be made publicly available through the MorPhiC Data Explorer (https://morphic.bio/data/), and are available from the authors on reasonable request in the interim, subject to MorPhiC consortium data-sharing policies. Benchmark comparison scripts and documentation are included in the repository under tests/ and docs/.

### Authors’ contributions

L.H.H. developed the software and wrote the manuscript. K.Y.Y. reviewed and contributed to the editing of the manuscript. D.B., B.F. and P.R. generated the JAX bulk RNA-seq and scRNA-seq (Flex) datasets. D.H., R.L. and T.Z. generated the MSK perturb-seq data.

### Competing interests

L.H.H. and K.Y.Y. have equity interest in Biodepot LLC. The terms of this arrangement have been reviewed and approved by the University of Washington in accordance with its policies governing outside work and financial conflicts of interest in research.

### Funding

L.H.H. and K.Y.Y. at University of Washington are supported by the National Institutes of Health (NIH) grant U24HG012674. K.Y.Y. is also supported by the Virginia and Prentice Bloedel Endowment at the University of Washington. D.B., B.F. and P.R. at The Jackson Laboratory are supported by NIH grant UM1HG012651. D.H., R.L. and T.Z. at the Memorial Sloan Kettering Cancer Center are supported by NIH grant UM1HG012654 and MSKCC Cancer Center Support Grant P30CA008748.

## Acknowledgements

We would like to thank the data ingestion team at the European Bioinformatics Institute (Anu Shivalikanjli, Galabina Yordanova, Alexandros Orges Koci, Robert Wilson and Helen Parkinson) for data collection, metadata annotation and brokering of the MorPhiC datasets. We would also like to thank the coordination team at the University of Miami (Dusica Vidovic and Stephan Schurer).

## Notes

https://github.com/morphic-bio/STAR-suite

https://github.com/morphic-bio/morphic-recipes

https://github.com/morphic-bio/morphic-provenance

### Summary of Updates

Flex probe assignment replaced by a disambiguated hash cascade that aligns nothing; all benchmarks re-run on release 1.9.5 against Cell Ranger 9.0.1, raising Flex speedups from 3.6-5.7-fold to 17-41-fold. Added comparison with cyto 0.4.7, three public benchmark datasets, and disk and memory measurements. Abstract restructured; supplementary expanded.

