## Supplemental Data 1 for "STAR Suite: Transcriptomics processing in a single binary through AI-assisted development"

Supplementary Table S1. scRNA-seq parameter comparison on three public datasets, each configuration against a Cell Ranger 9.0.1 run on the same FASTQ files (no BAM output, secondary analysis disabled): upstream STAR 2.7.11b with the community-optimized STARsolo parameters (CellGENI) and STAR Suite 1.9.5 in its Cell Ranger-compatible mode (poly-G trimming and a matched multimapper policy). Datasets: 10x PBMC 10K (3' v3, 638.9M read pairs); a glioma cell line (GEO GSE261042, SRA SRR28247645; 3' v3, 117.0M read pairs, about 18,000 reads per cell); an organoid (GEO GSE316010, SRA SRR36749062; GEM-X 3' v4, 301.6M read pairs). Spearman and Pearson are gene-level correlations of raw per-gene totals over the cells both tools called, over every gene in both annotations; Jaccard is on the called-cell sets; Cell Ranger 9.0.1 is the reference for both. Wall time is whole-process wall-clock time on the lab server (32 threads, every file on NVMe).

| Dataset | Configuration | Cells | Spearman | Pearson | Jaccard | Wall time |
|---|---|---|---|---|---|---|
| PBMC 10K | CellGENI (STAR 2.7.11b) | 11,794 | 0.950 | 0.993 | 0.994 | 30 min 43 s |
| | STAR Suite 1.9.5 | 11,863 | 0.990 | 0.9998 | 0.995 | 16 min 18 s |
| | Cell Ranger 9.0.1 | 11,806 | — | — | — | 63 min 0 s |
| GSE261042 | CellGENI (STAR 2.7.11b) | 6,430 | 0.957 | 0.997 | 0.998 | 8 min 0 s |
| | STAR Suite 1.9.5 | 6,441 | 0.992 | 0.9993 | 0.998 | 5 min 12 s |
| | Cell Ranger 9.0.1 | 6,427 | — | — | — | 19 min 31 s |
| GSE316010 | CellGENI (STAR 2.7.11b) | 10,353 | 0.960 | 0.999 | 0.968 | 17 min 36 s |
| | STAR Suite 1.9.5 | 10,352 | 0.993 | 0.99994 | 0.971 | 11 min 27 s |
| | Cell Ranger 9.0.1 | 10,060 | — | — | — | 38 min 34 s |

Supplementary Table S2. STAR-Perturb runtime and CRISPR-call concordance across two Perturb-seq benchmarks (STAR Suite 1.9.5; Cell Ranger 9.0.1 without BAM output or secondary analysis; Intel i9-13900KF, 32 threads, all files on NVMe). The MSK Cell Ranger runtime is the sum of the separate GEX + gRNA and GEX + LARRY runs. CR match, fraction of shared cells with identical guide calls; UMI corr., per-guide UMI-sum Pearson over shared cells. CR, Cell Ranger.

| Dataset | STAR | CR | Speedup | CR match | UMI corr. |
|---|---|---|---|---|---|
| A375 | 2 min 26 s | 11 min 25 s | 4.7× | 100% | 0.99999 |
| MSK 30-KO | 27 min 6 s | 167 min 2 s | 6.2× | 99.0% | 0.9994 |

### Supplementary Note 1: measured comparison with cyto on the three Flex benchmarks

cyto (Teyssier and Dobin, bioRxiv 2026) is an open-source Flex processor based on k-mer lookup without alignment. We ran cyto 0.4.7 on all three Flex benchmarks of Table 1 of the main text: the JAX SC2300771 and GSE325982 datasets on the same 32-thread machine used for the other benchmarks in this paper, and the 320k dataset on the same AWS instance as its Cell Ranger and STAR-Flex runs (Methods), under cyto's preferred conditions: BINSEQ (CBQ) input as well as gzipped FASTQ, read geometry confirmed by cyto's own detection tool, the probe-barcode list

Supplementary Table S3. Concordance of STAR-Flex with Cell Ranger 9.0.1, sample by sample and pooled, on the three multiplexed Flex datasets: JAX SC2300771 (four biological samples, one tag each; our own Cell Ranger run), the public GSE325982 pool (four single-tag samples; our own Cell Ranger run, which calls the same cells as the submitters' deposited outputs with every concordance measure at 1.000) and the 10x 320k scFFPE dataset (eight samples, two tags each; our Cell Ranger run on the AWS instance). A cell is identified by its 16-base barcode together with its sample tag, since cells from different samples can share a barcode. Barcode Jaccard is computed on the called-cell sets. Cell Pearson correlates each shared cell's total count across cells. Mean per-cell Pearson correlates each shared cell's log-transformed counts across genes with at least 20 counts in both outputs and detected in at least 1% of shared cells, averaged over cells. Gene Spearman and gene Pearson are on raw per-gene totals over shared cells, over every probe-set gene in both outputs (Methods). "Mean over samples" averages the per-sample values and so also reflects any disagreement in assigning reads to samples; "All samples pooled" computes each measure once over all samples together and gives the values quoted in the main text. Cell counts in both summary rows are summed over samples. Computed on the STAR Suite 1.9.5 outputs (for the 320k dataset, the 32-thread run with in-memory buckets) with the concordance script in the benchmark repository. CR, Cell Ranger.

| Sample (tags) | STAR cells | CR9 cells | Jaccard | Cell Pearson | Mean per-cell Pearson | Gene Spearman | Gene Pearson |
|---|---|---|---|---|---|---|---|
| *JAX SC2300771* | | | | | | | |
| WT-Day-7 (BC004) | 4,533 | 4,397 | 0.970 | 0.999983 | 0.9981 | 0.9999631 | 0.9999990 |
| PAX6-PTC-D9-Day7 (BC006) | 5,417 | 5,343 | 0.986 | 0.999984 | 0.9987 | 0.9999701 | 0.9999990 |
| WT-Day-8 (BC007) | 5,399 | 5,383 | 0.997 | 0.999987 | 0.9987 | 0.9999568 | 0.9999995 |
| PAX6-PTC-D9-Day8 (BC008) | 5,308 | 5,296 | 0.997 | 0.999986 | 0.9988 | 0.9999670 | 0.9999995 |
| Mean over samples | 20,657 | 20,419 | 0.987 | 0.999985 | 0.9986 | 0.9999643 | 0.9999992 |
| **All samples pooled** | **20,657** | **20,419** | **0.988** | **0.999985** | **0.9986** | **0.9999746** | **0.9999997** |
| *GSE325982* | | | | | | | |
| OS01 (BC001) | 11,038 | 10,979 | 0.994 | 0.999969 | 0.9954 | 0.9999129 | 0.9999993 |
| OS02 (BC002) | 10,323 | 10,260 | 0.994 | 0.999968 | 0.9954 | 0.9999400 | 0.9999989 |
| OS03 (BC003) | 9,893 | 9,836 | 0.994 | 0.999972 | 0.9950 | 0.9998998 | 0.9999994 |
| OS04 (BC004) | 7,417 | 7,369 | 0.994 | 0.999975 | 0.9949 | 0.9999070 | 0.9999988 |
| Mean over samples | 38,671 | 38,444 | 0.994 | 0.999971 | 0.9952 | 0.9999149 | 0.9999991 |
| **All samples pooled** | **38,671** | **38,444** | **0.994** | **0.999971** | **0.9952** | **0.9999616** | **0.9999993** |
| *10x 320k scFFPE* | | | | | | | |
| BreastCancer1 (BC007–008) | 15,594 | 15,134 | 0.967 | 0.999996 | 0.9992 | 0.9999778 | 0.9999990 |
| Colorectal (BC003–004) | 54,062 | 52,560 | 0.971 | 0.999982 | 0.9990 | 0.9999804 | 0.9999998 |
| Endo (BC015–016) | 45,643 | 44,550 | 0.975 | 0.999988 | 0.9991 | 0.9999863 | 0.9999988 |
| Glioblastoma (BC001–002) | 34,105 | 33,342 | 0.974 | 0.999993 | 0.9990 | 0.9999843 | 0.9999983 |
| Kidney (BC011–012) | 34,421 | 33,687 | 0.976 | 0.999995 | 0.9991 | 0.9999893 | 0.9999997 |
| LNReactive (BC009–010) | 32,891 | 31,668 | 0.958 | 0.999994 | 0.9991 | 0.9999675 | 0.9999983 |
| LungCancer2 (BC005–006) | 70,395 | 69,330 | 0.984 | 0.999978 | 0.9990 | 0.9999848 | 0.9999998 |
| SkinMelanoma (BC013–014) | 46,300 | 45,139 | 0.972 | 0.999988 | 0.9990 | 0.9999858 | 0.9999997 |
| Mean over samples | 333,411 | 325,410 | 0.972 | 0.999989 | 0.9991 | 0.9999820 | 0.9999992 |
| **All samples pooled** | **333,411** | **325,410** | **0.974** | **0.999980** | **0.9990** | **0.9999997** | **0.9999997** |

Supplementary Table S4. STAR-SLAM parity with GRAND-SLAM on 100K-read human benchmark.

| SNP Mode | Threshold | Genes | NTR Pearson | NTR Spearman | k/nT Pearson |
|---|---|---|---|---|---|
| External BED mask | ≥20 reads | 384 | 0.9989 | 0.9810 | 0.9999 |
| External BED mask | ≥50 reads | 76 | 0.9961 | 0.9867 | 0.9984 |
| External BED mask | ≥100 reads | 23 | 0.9944 | 0.9941 | 0.9968 |

Supplementary Table S5. Benchmark runtimes and speedups, STAR Suite 1.9.5. Baseline is Cell Ranger 9.0.1 (no BAM output, secondary analysis disabled, whole-process wall time to completion) except bulk RNA-seq, whose baseline is an external stepwise pipeline (Trim Galore + STAR + Salmon); the with-Y-removal bulk baseline adds an external `awk`/`samtools`/`gzip` Y-removal step. Bulk, scRNA-seq, Perturb-seq and SLAM-seq rows ran on an Intel i9-13900KF at 32 threads with every file on local NVMe. The 320k Flex rows ran for STAR-Flex and Cell Ranger alike on an AWS instance with local NVMe, limited to the same size (m6id.16xlarge at 32 virtual CPUs and 128 GiB). Times are whole-process wall-clock seconds from `/usr/bin/time`. For reference, Cell Ranger writes its per-sample matrices before the pipeline completes (times taken from its run logs and output timestamps): at 3,054 s of the JAX run, 3,668 s of the GSE325982 run, 3,658 s of the PBMC run and 575 s of the A375 run. The speedups above use the completion time, since the output directory is delivered only then. The SLAM-seq row has no baseline: GRAND-SLAM runs on STAR's alignments and handles known variants differently from STAR-SLAM, so no like-for-like timing exists (main text, Results). Of the rows that read FASTQ, the PBMC, A375 and original-BGZF Flex rows are read by the in-process parallel BGZF reader; the MSK files are ordinary gzip and were read through STAR's internal gzip route, and the bulk and SLAM-seq runs use STAR's external `zcat` route, as their production commands do. For the modules that align, the route makes little difference to the time (main text, Results).

| Assay | Benchmark | STAR Suite (s) | Baseline (s) | Speedup |
|---|---|---|---|---|
| Bulk RNA-seq | PPARG 35.1M PE, no Y-removal | 485.8 | 769.0 | 1.6× |
| Bulk RNA-seq | PPARG 35.1M PE, with Y-removal | 574.5 | 2,446.6 | 4.3× |
| scRNA-seq | 10x PBMC 10K, 3' GEX | 977.8 | 3,780.0 | 3.9× |
| Perturb-seq | A375, GEX + CRISPR | 146.2 | 684.9 | 4.7× |
| Perturb-seq | MSK 30-KO ES, GEX + guides + LARRY (one pass vs. two CR runs) | 1,625.9 | 10,022.2 | 6.2× |
| 10x Flex | JAX SC2300771, original BGZF FASTQ | 199.8 | 3,426.1 | 17.2× |
| 10x Flex | JAX SC2300771, CBQ† | 169.8 | 3,426.1 | 20.2× |
| 10x Flex | GSE325982, plain gzip FASTQ via rapidgzip | 257.4 | 4,446.0 | 17.3× |
| 10x Flex | GSE325982, CBQ† | 108.4 | 4,446.0 | 41.0× |
| 10x Flex | 10x 320k scFFPE, original BGZF FASTQ | 1,055.9 | 23,858.5 | 22.6× |
| 10x Flex | 10x 320k scFFPE, CBQ† | 645.2 | 23,858.5 | 37.0× |
| SLAM-seq | GRAND-SLAM 100K fixture, external BED mask | 26.3 | — | — |

†Experimental indexed-binary (CBQ/BINSEQ) input path; the conversion is not included in the time.

restricted to the tags present in each dataset, and `uv` installed so that cyto's cell-filtering step ran as designed. In every comparison both tools read the same input files from the same device, with the page cache dropped before each timed run on an otherwise idle machine. Both also rely on reference-derived lookup structures that are excluded from neither timing in any material way: cyto builds its probe, whitelist and gene hashes at startup in 3.2 s, while STAR-Flex maps a prebuilt half-probe cache read-only at startup: an exact whole-probe table and two 25-base tables holding 8,296,122 single-mismatch half keys with the probes each can come from. Building that cache is a separate one-time step over the probe set, not the reads—2 min 18 s on 32 threads—and it is reused by every run against that probe set, so it is excluded from the per-run timings reported here. Timings used `/usr/bin/time -v`; the STAR-Flex and Cell Ranger figures are the recorded benchmarks of Supplementary Table S5. Concordance was computed against the same Cell Ranger 9.0.1 per-sample outputs for both tools, with the measures and the script of Supplementary Table S3: Jaccard on the called-cell sets, per-cell Pearson and Spearman correlations over the cells both tools called, with a cell identified by its barcode and sample tag and all samples pooled. cyto's values are computed from its filtered outputs; because cyto's cell calls vary by tens of cells between runs and between input formats (below), the run used is stated for each dataset: the storage-panel FASTQ

Supplementary Table S6. Variance classes: sources of residual difference between STAR Suite and the reference tools. The bracketed indices in the main-text parity table refer to these classes.

| Class | Source of residual difference |
|---|---|
| 1 | Stochastic filtering and random number generator (RNG) state (EmptyDrops Monte Carlo). The Monte Carlo draws are partitioned across bootstrap streams, so the stream count is part of the reproducibility contract: matched `-soloCellFilterBootstrapThreads` reproduces a callset exactly, while changing it moves a small number of marginal barcodes across the FDR boundary in both directions. Re-running the 320k benchmark with 32 streams where the reference used 48 shifted 28 of 333,439 called cells (0.008%; an earlier release, whose run called 333,439 cells against the 333,411 of the release benchmarked here), with every read-level counter unchanged |
| 2 | Marginal cell calls at the filtering boundary |
| 3 | Tie resolution—the order in which equivalent feature, alignment, or UMI candidates are accepted (a cross-cutting cause; see the main-text section on clean-room reimplementation and residual differences) |
| 4 | Partial sampling state, as in Salmon's dynamic library-type auto-detection where running totals are updated during sampling |
| 5 | Multithreaded read-order differences (bulk TranscriptVB residuals; matched single-threaded runs reproduce Salmon to within numerical noise) |
| 6 | Floating-point implementation differences during GRAND-SLAM EM replication (slightly different digamma-function implementations across libraries) |
| 7 | Ambiguous alignments (intronic and lncRNA regions) |
| 8 | Overlapping or poorly annotated genes |
| 9 | Inherent transcript-level ambiguity in isoform-level VB/EM |
| 10 | Version-dependent defaults (e.g., the Cell Ranger 7 default that counts intronic reads toward gene-level totals, which prevents direct STARsolo/Cell Ranger 7+ parity) |
| 11 | Clean-room black-box behavior with a documentation gap—behavior inferred from a proprietary reference's observable outputs alone rather than derived from its code (Cell Ranger Flex is the canonical case) |
| 12 | Intentional documented design choices (Hamming scoring of the unanchored probe half in Flex; connected-component UMI correction of feature barcodes in Perturb-seq) |

run for JAX, the laboratory-server FASTQ run for GSE325982, and an original-FASTQ run at 32 threads on the laboratory server for 320k. As a cross-check, the complete filtered output of the timed 32-thread cyto run on the 320k instance agrees with that local run at a called-cell Jaccard of 0.998 (345,528 shared cells; 280 and 310 called by one run only) and gives the same Jaccard against Cell Ranger, 0.883, with the same per-sample pattern.

**What each tool produces, relative to the comparator.** The comparator for every comparison in this paper is Cell Ranger's deliverable: per-sample filtered count matrices in MEX format, produced after cell calling. STAR-Flex writes exactly that. cyto offers two output modes, and only one of them reaches the comparator. Its matrix-format mode (`-format mtx`) writes the raw count matrix and stops: the cell-calling step is not run, so empty droplets remain in the matrix and no filtered matrix is produced. According to its documentation, cell calling is available only in its default h5ad mode, where it is performed by separate post-processing steps after counting: the raw matrix is converted to h5ad by one Python tool and filtered by another, both invoked by the main program. That conversion has no counterpart in STAR-Flex or Cell Ranger. We therefore benchmark cyto in h5ad mode, the only configuration that performs the same work, and report its wall time as measured, including the conversion. cyto's own per-module timing table attributes the conversion to 6–9 s per sample on the JAX benchmark, most of it overlapped with the processing of other samples. In the matrix-format mode that would match our output format, cyto does less work than the comparator requires.

**Cell calling on multiplexed samples.** cyto filters each tag separately and writes one filtered matrix per tag; a sample spread over several tags is assembled afterwards by concatenating those per-tag outputs, which is what its aggregation tool documents (`https://github.com/ArcInstitute/pycyto#aggregate`) and what its maintainer recommends when asked (`https://github.com/ArcInstitute/cyto/issues/238#issuecomment-4170963800`); cyto 0.4.7 exposes no option that fits one cell-calling model across a sample's tags. Our comparison follows that route, concatenating the per-tag filtered outputs by sample with tag-qualified cell identities. Cell Ranger calls cells per biological sample, pooling the tags that belong to one sample into one model, and STAR-Flex does the same (Methods, Cell calling); in the pooled model the thresholds and the ambient profile are estimated once, on the whole sample. The two procedures differ, and the agreement measured below differs with them; attributing the whole of the difference to per-tag filtering would need a controlled comparison that we have not run. On the JAX SC2300771 and GSE325982 datasets every sample is a single tag, so the distinction does not arise and the three tools agree closely (Table S7). On the 320k dataset each of the eight samples is spread over two tags, and there cyto's agreement with Cell Ranger's called cells falls to 0.696–0.952 per sample (pooled 0.883), against 0.958–0.984 (pooled 0.974) for STAR-Flex (Table S8); the per-cell expression correlations on the cells both tools call remain high for both. Because each tool is otherwise scored on its own set of shared cells, we also compared the two on the cells all three tools call (Table S8, lower panel): on those identical 314,388 cells STAR-Flex agrees with Cell Ranger at a mean per-cell Pearson of 0.9991 and Spearman of 0.9993, against 0.9958 and 0.9948 for cyto. The 320k dataset is the largest public Flex benchmark and is multiplexed the way the consortium's own data are.

**Chip-generation check for the 320k GEM-X dataset.** Before benchmarking, we confirmed that the GEM-X 320k dataset presents the same inputs to processing software as the Next GEM JAX SC2300771 dataset. Sampling 100,000 raw read pairs from lane 1 of the 320k FASTQ files: read 1 is 28 bases (16-base cell barcode followed by a 12-base UMI), and 97.1% of reads carry a cell barcode found exactly—before any error correction—in the 737,280-entry `737K-fixed-rna-profiling` whitelist (the JAX SC2300771 Next GEM dataset measures 90.9% on the same test); read 2 is 90 bases, and the 8-base sample barcode at positions 69–76 matches Cell Ranger's BC001–BC016 recognition table in 94.9% of reads; the dataset's published configuration specifies probe set v1.1.0 on the GRCh38-2024-A reference, identical to the JAX benchmark stack. Cell Ranger 9.0.1 provides a single cell-barcode whitelist for Flex, used for both chip generations.

On the public 320k scFFPE GEM-X Flex dataset (Table 1 of the main text), STAR-Flex 1.9.5 completes in 17 min 36 s from the delivered BGZF files and 10 min 45 s from CBQ input at 32 threads with 72–74 GiB peak memory—the cell-barcode buckets held entirely in memory—and called-cell agreement with the Cell Ranger 9 reference outputs of Jaccard 0.974 (333,411 cells against 325,410; 325,036 shared) gene Spearman 0.99998 and gene Pearson 0.999999, averaged over the eight samples. cyto reports 13 min on the same dataset at 128 threads from BINSEQ input; measured on the same instance at 32 threads from the same original files, cyto 0.4.7 completed in 23 min 16 s (14 min 48 s from CBQ), so STAR-Flex is 1.32-fold faster on identical bytes (1.38-fold from CBQ), and Cell Ranger 9.0.1 took 6 h 37 min 38 s (Table S7); the cell-calling comparison on this multiplexed dataset is given above (Table S8). On the spinning-disk instance (Supplementary Table S9) the same runs took 30 min 5 s for STAR-Flex, 1 h 29 min 1 s for cyto and 9 h 0 min 11 s for Cell Ranger. Raw per-cell and per-gene comparison tables and the scripts that produce them are provided with the benchmark repository.

Supplementary Table S7. STAR-Flex 1.9.5 and cyto 0.4.7 on the three Flex benchmarks, each against the same Cell Ranger 9.0.1 per-sample outputs; the concordance rows compare each tool with Cell Ranger rather than with one another, while the time and memory rows are a direct comparison on the same hardware. JAX and GSE325982 were measured on the laboratory server (32 threads, local NVMe) and 320k on an AWS m6id.16xlarge limited to the same size (Methods). Concordance pools each dataset's samples with a cell identified by barcode and tag (Supplementary Table S3); cyto's cell calls are from the run named in the text; the calls of its timed runs agree with them at Jaccard ≥0.998. STAR-Flex memory on 320k is with the cell-barcode buckets held in memory.

| | **STAR-Flex** | **cyto** | **Cell Ranger 9.0.1** |
|---|---|---|---|
| *JAX SC2300771 (Next GEM; 4 single-tag samples; 2.011B read pairs)* | | | |
| Wall time, original BGZF FASTQ | 3 min 20 s | 3 min 40 s | 57 min 6 s |
| Wall time, CBQ/BINSEQ | 2 min 50 s | 3 min 36 s | — |
| Peak resident memory | 24 GiB | 38 GiB | — |
| Called-cell Jaccard vs. Cell Ranger | 0.988 | 0.985 | (reference) |
| Mean per-cell Spearman | 0.9988 | 0.9946 | (reference) |
| Mean per-cell Pearson | 0.9986 | 0.9953 | (reference) |
| *GSE325982 (Next GEM; 4 single-tag samples; 1.121B read pairs)* | | | |
| Wall time, plain gzip FASTQ | 4 min 17 s | 5 min 25 s | 74 min 6 s |
| Wall time, CBQ/BINSEQ | 1 min 48 s | 2 min 14 s | — |
| Peak resident memory | 18 GiB | 37 GiB | — |
| Called-cell Jaccard vs. Cell Ranger | 0.9939 | 0.9933 | (reference) |
| Mean per-cell Spearman | 0.9931 | 0.9910 | (reference) |
| Mean per-cell Pearson | 0.9952 | 0.9940 | (reference) |
| *10x 320k scFFPE (GEM-X; 8 samples on 16 tags, two tags per sample; 7.30B read pairs)* | | | |
| Wall time, original BGZF FASTQ | 17 min 36 s | 23 min 16 s | 6 h 37 min 38 s |
| Wall time, CBQ/BINSEQ | 10 min 45 s | 14 min 48 s | — |
| Peak resident memory | 72 GiB | 115 GiB | — |
| Called-cell Jaccard vs. Cell Ranger | 0.974 | 0.883 | (reference) |
| Mean per-cell Spearman | 0.9992 | 0.9949 | (reference) |
| Mean per-cell Pearson | 0.9990 | 0.9958 | (reference) |

### Supplementary Note 2: Flex on spinning disk

Cell Ranger's Flex pipeline needs about 3 TB of fast scratch per concurrent run on the 320k dataset (main text), and sites that cannot provide it place their working files on spinning or network storage, where the volume a tool writes converts directly into elapsed time. We therefore repeated the JAX, GSE325982 and 320k benchmarks with every input, output, temporary and spill location on spinning disk, holding the tools, the data and every other setting fixed, and ran Cell Ranger 9.0.1 under the same placement (Supplementary Table S9).

So that the two placements differ in storage rather than processor, both were measured on AWS instances limited to the lab server's size—32 virtual CPUs and 128 GiB of memory, Ubuntu 24.04: the m6id.16xlarge used for the 320k speed comparison (Intel Xeon Platinum 8375C, two local NVMe devices in RAID0), and an h1.8xlarge (Intel Xeon E5-2686 v4) with four local hard drives in RAID5, the layout of our own array. Because the two instance types carry different processor generations, one statically linked compute-only program, a fixed integer recurrence with no input or output, was run on both after the benchmarks, pinned to 1, 16 and 32 virtual CPUs; the NVMe instance delivered 1.1, 1.4 and 1.8% more work per second. On both instances every tool ran at 32 threads, Cell Ranger with 32 cores and 120 GB of memory, and cyto under a 120 GiB memory limit.

Supplementary Table S8. Agreement with Cell Ranger 9.0.1 per sample on the 320k dataset, where each sample is multiplexed across two tags. Cells are identified by barcode and tag; STAR-Flex values in the upper panel are those of Supplementary Table S3. The lower panel compares expression on the triple intersection—the cells Cell Ranger, STAR-Flex and cyto all call—so that both tools are scored on identical cells (314,388 of Cell Ranger's 325,410). Pearson is the mean over cells of the correlation across genes of one cell's $\log(1+x)$ counts in the two outputs and Spearman the corresponding rank correlation, both as defined in Methods; the cells column sums over samples and the correlations are means over the eight samples. Gene-level totals over the same cells agree at Pearson $\geq$0.99999 and Spearman $\geq$0.99996 for both tools.

| | | **STAR-Flex** | | **cyto** | |
|---|---|---|---|---|---|
| **Sample** | **Cell Ranger cells** | **cells** | **Jaccard** | **cells** | **Jaccard** |
| Glioblastoma (BC001–002) | 33,342 | 34,105 | 0.974 | 34,835 | 0.923 |
| Colorectal (BC003–004) | 52,560 | 54,062 | 0.971 | 50,084 | 0.873 |
| Lung cancer (BC005–006) | 69,330 | 70,395 | 0.984 | 68,578 | 0.941 |
| Breast cancer (BC007–008) | 15,134 | 15,594 | 0.967 | 21,727 | 0.696 |
| Reactive lymph node (BC009–010) | 31,668 | 32,891 | 0.958 | 38,324 | 0.745 |
| Kidney (BC011–012) | 33,687 | 34,421 | 0.976 | 34,948 | 0.952 |
| Skin melanoma (BC013–014) | 45,139 | 46,300 | 0.972 | 47,590 | 0.924 |
| Endometrium (BC015–016) | 44,550 | 45,643 | 0.975 | 49,722 | 0.884 |
| Pooled | 325,410 | 333,411 | 0.974 | 345,808 | 0.883 |
| *Expression on the cells all three tools call* | | | | | |
| | **Cells** | **Pearson** | **Spearman** | **Pearson** | **Spearman** |
| Glioblastoma (BC001–002) | 32,690 | 0.9990 | 0.9995 | 0.9953 | 0.9947 |
| Colorectal (BC003–004) | 47,847 | 0.9990 | 0.9991 | 0.9957 | 0.9947 |
| Lung cancer (BC005–006) | 66,855 | 0.9990 | 0.9991 | 0.9961 | 0.9950 |
| Breast cancer (BC007–008) | 15,105 | 0.9992 | 0.9994 | 0.9959 | 0.9950 |
| Reactive lymph node (BC009–010) | 29,806 | 0.9991 | 0.9994 | 0.9960 | 0.9952 |
| Kidney (BC011–012) | 33,422 | 0.9991 | 0.9993 | 0.9957 | 0.9946 |
| Skin melanoma (BC013–014) | 44,463 | 0.9990 | 0.9993 | 0.9954 | 0.9945 |
| Endometrium (BC015–016) | 44,200 | 0.9991 | 0.9992 | 0.9960 | 0.9951 |
| All samples | 314,388 | 0.9991 | 0.9993 | 0.9958 | 0.9948 |

Against cyto the ordering is the same at both placements but the margin widens. Moving from NVMe to spinning disk costs STAR-Flex up to 2.1-fold and cyto up to 5.3-fold, so on spinning disk STAR-Flex is 1.5- to 3.0-fold faster than cyto from the original files and 2.3- to 3.9-fold from CBQ, against 1.1- to 1.4-fold on NVMe; on the 320k set it finishes in 30 min against 1 h 29 min. The cause is visible in the disk each tool needs for the same work—2.5 and 4.6 GiB against 76 and 39 GiB on the two smaller datasets, 16.9 GiB against 218 GiB on the 320k set—and in how far each falls short of the machine: on spinning disk STAR-Flex keeps 9 to 15 of the 32 virtual CPUs busy and cyto 3 to 5, against 25 to 29 and 12 to 18 on NVMe, so both become bound by the disk, but the tool that writes tens of times more waits proportionally longer for it. Against Cell Ranger the margin narrows, because Cell Ranger spends most of its run computing rather than waiting on the disk: it slows by 13 to 37%, taking 9 h on the 320k set, and STAR-Flex remains 12- to 20-fold faster from the original files and 16- to 37-fold from CBQ. Only the runs that read a single stream of plain gzip change little between placements—STAR-Flex and cyto on the original GSE325982 pair—because they are bound by decompressing that stream rather than by the disk.

Supplementary Table S9. Flex wall-clock time by storage placement. Both placements were measured on AWS instances limited to the lab server's size (32 virtual CPUs, 128 GiB) that differ in storage: m6id.16xlarge with two local NVMe devices in RAID0, and h1.8xlarge with four local hard drives in RAID5. A compute-only control delivered 1–2% more throughput on the NVMe instance (see text). Every input, output, temporary and spill location was on the stated device. STAR Suite 1.9.5 with cell-barcode buckets in memory; cyto 0.4.7 under a 120 GiB memory limit; Cell Ranger 9.0.1 with no BAM output and secondary analysis disabled, timed to completion; all at 32 threads, and every run completed in one attempt. Ratio is spinning-disk time over NVMe time. Peak disk is the high-water mark of filesystem use above the pre-run baseline, sampled every 5 s on the NVMe instance, which is what a site has to provision; a tool that deletes intermediates as it goes, as Cell Ranger does, occupies less at any moment than it writes over the run. On spinning disk the peaks were within 2% for STAR Suite and Cell Ranger and up to 17% higher for cyto. The NVMe times for JAX and GSE325982 differ from Table 2 of the main text, which was measured on the lab server; the speedups over Cell Ranger agree (16.0- and 17.9-fold from the original files here, 17.2- and 17.3-fold there).

| Dataset | Tool, input | NVMe | Spinning disk | Ratio | Peak disk |
|---|---|---|---|---|---|
| JAX SC2300771 | STAR-Flex, original BGZF | 5 min 14 s | 9 min 47 s | 1.87 | 2.5 GiB |
| | STAR-Flex, CBQ | 3 min 24 s | 7 min 17 s | 2.14 | 2.5 GiB |
| | cyto, original FASTQ | 5 min 48 s | 20 min 41 s | 3.57 | 75.9 GiB |
| | cyto, CBQ | 4 min 36 s | 18 min 10 s | 3.95 | 75.3 GiB |
| | Cell Ranger 9.0.1 | 1 h 24 min 0 s | 1 h 54 min 50 s | 1.37 | 564 GiB |
| GSE325982 | STAR-Flex, plain gzip via rapidgzip | 6 min 10 s | 6 min 24 s | 1.04 | 4.6 GiB |
| | STAR-Flex, CBQ | 2 min 20 s | 3 min 24 s | 1.45 | 4.6 GiB |
| | cyto, plain gzip | 8 min 23 s | 9 min 21 s | 1.12 | 39.0 GiB |
| | cyto, CBQ | 3 min 11 s | 7 min 50 s | 2.46 | 41.1 GiB |
| | Cell Ranger 9.0.1 | 1 h 50 min 15 s | 2 h 5 min 1 s | 1.13 | 254 GiB |
| 10x 320k scFFPE | STAR-Flex, original BGZF | 17 min 36 s | 30 min 5 s | 1.71 | 16.9 GiB |
| | STAR-Flex, CBQ | 10 min 45 s | 20 min 17 s | 1.89 | 16.9 GiB |
| | cyto, original FASTQ | 23 min 16 s | 1 h 29 min 0 s | 3.82 | 218 GiB |
| | cyto, CBQ | 14 min 48 s | 1 h 18 min 35 s | 5.31 | 208 GiB |
| | Cell Ranger 9.0.1 | 6 h 37 min 38 s | 9 h 0 min 11 s | 1.36 | 1,805 GiB |

### Supplementary Note 3: The alignment-validated cache rerun with the current cell caller

STAR-Flex's first single-process route assigned probes through an alignment-validated cache (main text). Its comparison with Cell Ranger predates the tag-aware cell caller, so to isolate the probe-assignment route we reran it with release 1.9.4 under the compatibility flag (`-flexLegacy yes`) with the legacy H0/H1 cache, as cache lookup only with no alignment, and with every other setting of the half-probe runs held fixed, on the JAX and 320k datasets. Both routes were then compared against the same Cell Ranger 9.0.1 outputs with the same script and measures as Supplementary Table S3 (Supplementary Table S10). On JAX the two routes are equivalent (pooled Jaccard 0.990 against 0.988). On 320k the alignment-validated cache falls to 0.933 pooled and 0.803 to 0.968 per sample, against 0.974 and 0.958 to 0.984 for the half-probe rule, and calls 318,331 cells against 333,411 (Cell Ranger 325,410). The two routes agree with each other at 0.989 on JAX and 0.945 on 320k. Expression agreement on the cells both outputs called stays above 0.994 for both routes on both datasets. The run logs show the cause. Reads that the validated cache cannot resolve are left unassigned and, with no alignment pass, discarded: about 8.2% of the JAX reads and 6.6% of the 320k reads, so the route keeps 8.6% and 4.8% fewer reads than the half-probe rule, whose seed-and-extend step resolves them. At the JAX depth a proportional loss moves few cells across the calling threshold; at the 320k depth it moves many. The runs and comparison are recorded in `analysis/flex_legacy_route_concordance_20260920`.

Supplementary Table S10. Concordance with Cell Ranger 9.0.1 of STAR-Flex's two probe-assignment routes, both run with release 1.9.4 and the same tag-aware cell caller. Top: pooled values, cells identified by barcode and tag, computed as in Supplementary Table S3; the cell counts in the first column are Cell Ranger's. Cell $r$, Pearson correlation of per-cell totals; per-cell $r$ and $\rho$, mean per-cell Pearson and Spearman; gene $\rho$ and $r$, gene Spearman and Pearson. Bottom: per-sample called-cell Jaccard on the 320k dataset. Alignment-validated: the legacy H0/H1 cache (`-flexLegacy yes`), cache lookup only, no alignment. Half-probe: the release default.

| **Dataset** | **Route** | **Cells** | **Jaccard** | **Cell $r$** | **Per-cell $r$** | **Per-cell $\rho$** | **Gene $\rho$** | **Gene $r$** |
|---|---|---|---|---|---|---|---|---|
| JAX (20,419) | align.-validated | 20,448 | 0.990 | 0.99998 | 0.9950 | 0.9946 | 0.9999460 | 0.9999409 |
| | half-probe | 20,657 | 0.988 | 0.99999 | 0.9986 | 0.9988 | 0.9999746 | 0.9999997 |
| 320k (325,410) | align.-validated | 318,331 | 0.933 | 0.99998 | 0.9962 | 0.9957 | 0.9999960 | 0.9999958 |
| | half-probe | 333,411 | 0.974 | 0.99998 | 0.9990 | 0.9992 | 0.9999997 | 0.9999997 |

| | | **Alignment-validated** | | **Half-probe** | |
|---|---|---|---|---|---|
| **320k sample** | **Cell Ranger cells** | **cells** | **Jaccard** | **cells** | **Jaccard** |
| Glioblastoma (BC001–002) | 33,342 | 34,001 | 0.968 | 34,105 | 0.974 |
| Colorectal (BC003–004) | 52,560 | 48,895 | 0.891 | 54,062 | 0.971 |
| Lung cancer (BC005–006) | 69,330 | 68,092 | 0.949 | 70,395 | 0.984 |
| Breast cancer (BC007–008) | 15,134 | 15,375 | 0.948 | 15,594 | 0.967 |
| Reactive lymph node (BC009–010) | 31,668 | 26,411 | 0.803 | 32,891 | 0.958 |
| Kidney (BC011–012) | 33,687 | 34,198 | 0.967 | 34,421 | 0.976 |
| Skin melanoma (BC013–014) | 45,139 | 46,045 | 0.962 | 46,300 | 0.972 |
| Endometrium (BC015–016) | 44,550 | 45,314 | 0.965 | 45,643 | 0.975 |
| Pooled | 325,410 | 318,331 | 0.933 | 333,411 | 0.974 |

### Supplementary Note 4: Cell Ranger probe filtering on GSE325982

Cell Ranger's probe set can be used in full or restricted to the probes it marks as included, and the included set is the default. Every run reported in this paper uses the default, on all three Flex datasets and for all three tools (Methods). GSE325982 is the one dataset whose submitters deposited outputs made the other way, with filtering off, so their matrices report 18,488 genes where the default reports 18,129.

To show what that setting changes, we ran Cell Ranger 9.0.1 on GSE325982 twice, once with the submitters' setting and once with the default, everything else equal. The run with their setting reproduces their deposited matrices byte for byte, which fixes the configuration. Against it, the default run calls exactly the same 38,444 cells, sample for sample, and reports the same counts on the genes both keep: per-cell and per-gene correlations are 1.000000 throughout (Table S11). What the default removes is 359 genes and 0.34% of the unique molecular identifiers, which is why the two runs' cell totals correlate at 0.9999998 rather than exactly 1.

This pair also serves as a control on the concordance measures used throughout the paper. Two runs of the same tool, differing only in that one setting, return 1.000000 on every per-cell and per-gene measure, so the measures have no noise floor of their own here: where a comparison with STAR-Flex or cyto falls below 1, the difference is in the outputs, not in the measurement.

Supplementary Table S11. Cell Ranger 9.0.1 on GSE325982 with the submitters' setting (probe filtering off; the configuration that reproduces their deposit byte for byte) against the default (included probes only), everything else equal. The default reports 18,129 genes against 18,488 and 606,327,232 unique molecular identifiers against 608,416,597 (99.66%). Cells are identified by barcode and tag. Correlations are over the genes both runs report, by the definitions of Methods; per-cell values are means over cells. Jaccard is on the called-cell sets.

| Sample | Cells (both) | Jaccard | Cell Pearson | Per-cell Pearson | Per-cell Spearman | Gene Spearman and Pearson |
|---|---|---|---|---|---|---|
| OS01 (BC001) | 10,979 | 1.000000 | 0.9999998 | 1.000000 | 1.000000 | 1.000000 |
| OS02 (BC002) | 10,260 | 1.000000 | 0.9999998 | 1.000000 | 1.000000 | 1.000000 |
| OS03 (BC003) | 9,836 | 1.000000 | 0.9999997 | 1.000000 | 1.000000 | 1.000000 |
| OS04 (BC004) | 7,369 | 1.000000 | 0.9999998 | 1.000000 | 1.000000 | 1.000000 |
| **All samples pooled** | **38,444** | **1.000000** | **0.9999998** | **1.000000** | **1.000000** | **1.000000** |
